# Legumain (asparaginyl endopeptidase) modulates extracellular matrix dynamics in pulmonary fibrosis

**DOI:** 10.64898/2026.07.20.739492

**Authors:** Baptiste Rigoux, Alexis David, Damien Sizaret, Rania Allouche, Clémentine Lebouché, Lise Vanderlynden, Fabien Lecaille, Marcin Poreba, Florian Veillard, Sylvain Marchand-Adam, Gilles Lalmanach, Ahlame Saidi

## Abstract

Pulmonary fibrosis is characterized by extracellular matrix (ECM) deposition driven by fibroblast-to-myofibroblast transition (FMT) and by an altered proteolytic balance. While the roles of several cysteine proteases have been documented, the specific contribution of cathepsin V (CatV) and legumain (LGMN) remains poorly explored. LGMN, CatV and their dual inhibitor cystatin M/E (CysM/E) are significantly increased in lung specimens and bronchoalveolar lavage fluids from patients with idiopathic pulmonary fibrosis. TGF-β1 triggered CysM/E expression and LGMN transcription, intracellular maturation, enzymatic activity, and pro-LGMN secretion via the Smad-3 pathway, whereas CatV was downregulated in human lung fibroblasts (CCD-19Lu and primary HPF cells) undergoing myodifferentiation. Genetic silencing of LGMN or CatV, and pharmacological inhibition of LGMN, led to accumulation of fibronectin and elastin, implying that both proteases contribute to ECM remodeling. LGMN cleaved fibronectin, while CatV predominantly regulated elastin levels. Conversely, broad-spectrum inhibitor cystatin C (hCC) markedly _r_educed elastin and fibronectin degradation, whereas CysM/E exerted a weaker effect, mainly on elastin turnover. LGMN inhibition transiently delayed fibroblast wound closure, establishing a functional role in tissue repair through fibronectin remodeling. Neither LGMN nor CatV influenced α-SMA expression, distinguishing them from CatB, which participates in FMT. Altogether, LGMN was identified as an effector of matrix remodeling rather than myodifferentiation, acting in concert with CatV. Within the proteolytic network governing fibrosis progression, the present findings identify a cystatin-regulated LGMN/CatV partnership, participating in ECM turnover and cell migration. Present results also provide new perspectives on potential therapeutic protease-based strategies targeting ECM turnover underlying lung fibrosis.

## INTRODUCTION

Pulmonary fibrosis (PF), including idiopathic pulmonary fibrosis (IPF) is a progressive and irreversible disease of unknown etiology, (median survival: 2-4 years) that is characterized by interstitial fibrotic thickening [1, 2]. During the early phase of the disease, alveolar epithelial cell injury triggers, via the release of TGF-β1, the proliferation of lung fibroblasts and their differentiation into myofibroblasts. ECM overproduction by myofibroblasts, which is closely associated with an altered protease/antiprotease balance, induces dysregulation of ECM remodeling in fibrotic foci [3]. Besides matrix metalloproteinases (MMPs), matriptase, human airway trypsin-like protease (HAT), as well as tissue inhibitors of MMPs (TIMPs) and plasminogen activator inhibitor (PAI)-1 [4, 5], increasing evidence supports the involvement of cysteine cathepsins in fibrogenesis and in the maintenance of pulmonary homeostasis [6]. Among eleven human members (i.e., cathepsins B, C, F, H, K, L, O, S, V, W, and X; clan CA, family C1), several cysteine cathepsins have been identified as potent collagenases and/or elastases that participate in the remodeling and degradation of ECM components [7–10]. In addition to its distinctive collagenolytic activity that restrains the deposition of type I collagen, cathepsin K (CatK) degrades the pro-fibrotic cytokine TGF-β1 [11] and has been proposed as a key regulator in lung homeostasis [12], whereas cathepsin B (CatB) promotes lung fibroblast myodifferentiation through activation of the canonical TGF-β1/Smad pathway [13]. Also, CatF and CatV (a.k.a., CatL2), which exhibit restricted tissue distribution, were reported as putative biomarkers of cardiac, liver and skin fibrosis (for review: [14]). Pharmacological targeting of CatB as well as CatS might be relevant in the treatment of fibrotic disorders (for review: [14–16]). On the other hand, cystatins (clan IH, family I25) are potent tight-binding inhibitors of cysteine cathepsins. Serum human cystatin C (hCC), a broad-spectrum cathepsin inhibitor, is recognized as valuable biomarker of cardiac and liver fibrosis (for review: [17]), while hCC of bronchoalveolar lavage fluids (BALFs) was proposed as IPF biomarker [18]. However, a contrasting result was established for its mouse counterpart, showing that murine cystatin C was not a reliable marker of fibrosis during experimental bleomycin-induced lung fibrosis [19]. hCC is oversecreted during TGF-β1-induced differentiation of lung fibroblasts, endorsing that TGF-β1 promotes fibrosis by driving the hCC-dependent inhibition of collagenolytic cathepsins [13, 20]. Besides type I collagen deposition, increased expression and deposition of both elastin and fibronectin in fibrotic foci are also observed during fibrogenesis. Related to this statement, CatV displays a potent elastinolytic activity [21, 22], while legumain (LGMN; clan CD, family C13) may contribute to ECM remodeling in various organs and modulate inflammatory and fibroblast-activating pathways, likely participating in fibronectin degradation [23, 24]. From a clinical perspective, high circulating LGMN (a.k.a., AEP: asparaginyl endopeptidase) levels correlate with improved progression-free survival in IPF patients, and LGMN was proposed as one of the six major biomarkers in an IPF clinical decision index [25]. Accordingly, this led us to consider the potential role of CatV and LGMN, given that their contribution to the pathogenesis of lung fibrosis and subsequent ECM remodeling remains largely unexplored. Of note, cystatin M/E (CysM/E; a.k.a., CST6: cystatin 6), a type 2 cystatin related to cystatin C, is a unique dual tight-binding inhibitor of both LGMN and cysteine cathepsins (especially CatV/L2 and CatL) via two distinct binding domains (for review: [26, 27]).

Here, we investigated the expression, regulation and functional contribution of LGMN and CatV in lung fibrosis using clinical specimens from patients with IPF and non-IPF controls (lung biopsies and bronchoalveolar lavage fluids (BALFs)) as well as two human lung fibroblast models: CCD-19Lu cells and primary human pulmonary fibroblasts (HPF). We also examined how the endogenous inhibitors - hCC and CysM/E - may influence the proteolytic balance governing fibronectin and elastin homeostasis. Attention was given to the role of the prevailing TGF-β1/Smad-3 signaling pathway driving FMT in the regulation of these proteases and their inhibitors. Collectively, our study provides some novel mechanistic clues related to a fibroblast LGMN-CatV axis, which could contribute to ECM remodeling and cell motility within the proteolytic network governing fibrosis progression.

## MATERIALS AND METHODS

### Human specimens

#### Cohort and ethics

Tumor-free lung tissues were obtained from patients undergoing surgery for lung tumors or diagnosed with idiopathic pulmonary fibrosis (IPF). Sample collection followed institutional guidelines. The study was approved by the local bioethics committee of the University Hospital Center (Tours, France) and both tissue and BALF collections were declared to the French Ministry of Higher Education, Research, and Innovation (approval DC-2008-308 for non-IPF biopsies, approval 2020/89196.1 for IPF biopsies, and approval DC-2010-1216 for BALF). Written informed consent was obtained from each patient in compliance with the Declaration of Helsinki. Ethical standards set out in the Declaration of Helsinki were respected.

#### Lung biopsies

Lung biopsies (27 non-IPF; 26 IPF) were placed in precooled Precellys tubes (2.8-mm ceramic beads), grindered and homogenized using a Precellys 24 tissue homogenizer (Bertin Technologies, Montigny-le-Bretonneux, France) in lysis buffer 1 (0.1 M sodium acetate buffer, pH 5.5, 0.04 mM pepstatin A, 0.5 mM EDTA, 0.5 mM AEBSF, 1 mM MMTS), according to the supplier’s protocol. Homogenates were centrifuged for 15 min (4 °C, 20,800 g) to remove tissue debris. Supernatants were collected and protein concentrations were determined by Bradford assay (Bio-Rad, Hercules, CA, USA).

#### BALFs

BALFs (10 non-IPF; 18 IPF) were collected under sterile conditions, processed and frozen at −80 °C, as described earlier [28]. BALFs were concentrated 20-fold, and protein concentrations werre measured (Bradford assay) prior to analysis.

### Recombinant proteins, chemicals, substrates, and inhibitors

Recombinant human LGMN, CatV, and TGF-β1 were purchased from R&D Systems Europe (Abingdon, UK). Recombinant human fibronectin was from Corning (Corning, NY, USA). Benzyloxycarbonyl-Arg-Arg-4-methylcoumarin-7-ylamide (Z-Arg-Arg-AMC) was obtained from Bachem (Bubendorf, Switzerland). Benzyloxycarbonyl-Phe-Arg-7-amido-4-methylcoumarin (Z-Phe-Arg-AMC), ethylenediaminetetraacetic acid (EDTA), S-methyl methanethiosulfonate (MMTS), N,N-dimethylformamide (DMF), and propidium iodide (PI) were purchased from Sigma-Aldrich (Saint-Quentin-Fallavier, France). DQ elastin was from Invitrogen (Waltham, MA, USA). Dithiothreitol (DTT) was obtained from Euromedex (Souffelweyersheim, France). L-3-carboxy-trans-2,3-epoxypropionyl-leucylamide-(4-guanido)-butane (E-64), pepstatin A, 4-(2-aminoethyl)benzenesulfonyl fluoride hydrochloride (AEBSF), ethyl (E)-4-(1-(2-amino-2-oxoethyl)-2-(((benzyloxy)carbonyl)-L-alanyl-L-alanyl)-hydrazinyl)-4-oxobut-2-enoate (RR-11a), BV6 (an antagonist of cIAP1 and XIAP) and staurosporine were from MedChemExpress (Sollentuna, Sweden). 6,7-Dimethoxy-2-((2E)-3-(1-methyl-2-phenyl-1H-pyrrolo[2,3-b]pyridin-3-yl)-prop-2-enoyl))-1,2,3,4-tetrahydroisoquinoline (SIS3) was obtained from Calbiochem (VWR International, Pessac, France). Morpholinourea-leucinyl-homophenylalanine-vinyl-sulfone phenyl (LHVS) was a kind gift from Prof. James H McKerrow (Skaggs School of Pharmacy and Pharmaceutical Sciences, University of California, San Diego, CA, USA). Ac-dTyr-Tic-Ser-Asn-ACC (dY-ACC, LGMN substrate) and Biotin-Ahx-dTyr-Tic-Ser-Asp-AOMK (MP-L01, LGMN inhibitor) were synthesized as previously reported [29].

### Western blotting

Antibody dilutions and sources were as follows: mouse anti-α-smooth muscle actin (α-SMA) (1:1000) and mouse anti-β-actin (1:1000) were purchased from Sigma-Aldrich. Mouse anti-human glyceraldehyde-3-phosphate dehydrogenase (GAPDH) (1:10,000) was purchased from Proteintech (Rosemont, IL, USA). Rabbit anti-human fibronectin (1:1000) was from Cell Signaling Technology (Danvers, MA, USA), and mouse anti-tropoelastin (1:500) was from Santa Cruz Biotechnology (Heidelberg, Germany). Goat anti-human LGMN (1:500), and mouse anti-human mature CatV (1:500) were obtained from R&D Systems. Rabbit anti-human pro-CatV (1:1000) was from Abcam (Cambridge, UK). Goat anti-human CatB and CatL (1:800) were from R&D Systems. Goat anti-human CysM/E (1:300), and mouse anti-hCC (1:300) were from R&D Systems. Goat anti-rabbit IgG-peroxidase conjugate, goat anti-mouse IgG-peroxidase conjugate and rabbit anti-goat IgG-peroxidase conjugate were from Sigma-Aldrich.

Cell supernatants were harvested in 0.1 M sodium acetate buffer, pH 5.5, containing protease inhibitors (0.5 mM AEBSF, 0.5 mM EDTA, 1 mM MMTS and 0.04 mM pepstatin A). Supernatants were concentrated 50-fold and centrifuged for 15 min (4 °C, 20,800 g) to remove debris. Cell layers were washed with ice-cold PBS and harvested by scraping in ice-cold 0.1 M sodium acetate buffer, pH 5.5, 0.5 mM AEBSF, 0.5 mM EDTA, 1 mM MMTS and 0.04 mM pepstatin A, supplemented with 0.5 % NP-40. Protein extraction was performed using three freeze-thaw cycles with liquid nitrogen followed by sonication for 2 min (8 cycles of 15 sec ON – 10 sec OFF). Lysates were then centrifuged for 15 min (4 °C, 20,800 g) and protein concentration was determined using a Bradford assay. Samples were mixed with Laemmli buffer and boiled for 5 min. Concentrated culture media (10 µg/well) and cell layer lysates (20 - 50 µg/well) were loaded onto SDS-PAGE gels (for analyses of fibronectin and elastin (10 % SDS-PAGE), α-SMA and proteases (12 % SDS-PAGE) and cystatins (15 % SDS-PAGE), respectively) and electrophoresis was carried out under reducing conditions. Prestained molecular weight standards, Precision Plus Protein Dual Color and Dual Xtra Standards, were obtained from Bio-Rad. Separated proteins were transferred onto a nitrocellulose membrane (Amersham Biosciences, Buckinghamshire, UK), which was blocked with 5 % nonfat powdered milk in PBS 0.1 % Tween-20. Membranes were incubated with primary antibodies overnight at 4 °C with agitation, followed by incubation with secondary antibodies (1:5000) for 2 h at room temperature. Proteins were detected by chemiluminescence using the ECL Plus or ECL Prime Western Blotting Detection System (Amersham Biosciences) according to manufacturer’s instructions. Equal protein loading was monitored using anti-β-actin or anti-GAPDH antibodies. Alternatively, protein loading was verified by Ponceau S staining (Sigma-Aldrich). Bands were quantified by densitometric analysis using the ImageJ software (NIH, Bethesda, MD, USA).

### Protease activity assays

#### LGMN activity

LGMN peptidase activity in cell lysates and biopsy extracts was measured at 37 °C using dY-ACC (20 µM) as substrate (λex = 350 nm; λem = 460 nm). Kinetic measurements were performed in 50 mM 2-(N-morpholino) ethanesulfonic acid buffer (MES) pH 5.0 with 250 mM NaCl, 5 mM DTT and 0.01 % Brij35 (polyethylene glycol lauryl ether, Thermo Fisher Scientific, Villebon-sur-Yvette, France).

#### Cysteine cathepsins

Cathepsin peptidase activity in concentrated conditioned media was measured at 37 °C using Z-Phe-Arg-AMC (20 μM) as substrate (λ-ex = 350 nm; λ-em = 460 nm). CatB-specific activity in cell lysates was measured at 37 °C using Z-Arg-Arg-AMC (20 μM) as substrate (λex = 350 nm; λem = 460 nm). Both kinetics were performed in 0.1 M sodium acetate buffer, pH 5.5 with 5 mM DTT and 0.01 % Brij35. All assays were conducted in 96-well plates (Nunc, Thermo Fisher Scientific) using a SpectraMax Gemini M2 microplate reader (Molecular Devices, Wokingham, Berkshire, UK). Control experiments were performed in the presence of MP-L01, E-64, or CA-074 (10 µM) as indicated.

#### Elastinolytic activity

CatV-dependent elastolysis was measured in cell lysates at 37°C using DQ-elastin as substrate (15 µg) in cathepsins activity buffer. Assays were performed in the presence of a specific CatS inhibitor, LHVS (10 nM). Fluorescence (λ-exc = 495 nm; λ-em = 555 nm) was recorded on a Varian Cary Eclipse spectrofluorometer (Agilent Technologies, Les Ulis, France). Control experiments were conducted in the presence of E-64 (10 µM).

#### Fibronectin degradation assays

Recombinant fibronectin (1 µM) was incubated with activated CatV or LGMN (enzyme:substrate ratios 1:100 to 1:10,000) for 6 h at 37 °C in their respective activity buffers (CatV buffer: 0.1 M sodium acetate buffer, pH 5.5 with 1 mM EDTA, 2 mM DTT and 0.01 % Brij 35; LGMN buffer: 0.1 M sodium acetate buffer, pH 5.5 with 1 mM EDTA, 2 mM DTT and 0.01 % Brij 35). Control experiments were carried out in the presence of E-64 (for CatV) or RR-11a (for LGMN) at 10 µM. Reactions were stopped by the addition of reducing Laemmli buffer and analyzed by SDS-PAGE (10 %) followed by Western blotting with an anti-fibronectin antibody, as described above.

### ELISA, soluble collagen quantification

Concentrations of LGMN and hCC in culture media or tissue extracts were determined using DuoSet sandwich ELISA kits (R&D Systems), according to the supplier’s recommendations. CysM/E concentration was determined using a sandwich ELISA kit provided by Invitrogen. CatV concentration was measured using a sandwich ELISA kit from Novus Biologicals. Absorbance was measured at 450 nm (SpectraMax ABS Plus Microplate Reader, Molecular Devices). Soluble collagen in the culture media was quantified using the Sircol collagen assay (Biocolor Ltd., Belfast, UK). Sirius red dye (1 mL) was added to concentrated cell supernatants and incubated for 30 min with gentle mixing. The collagen-dye complex was precipitated by centrifugation (10,000 g) for 10 min and dissolved in 0.5 M NaOH. Absorbance was determined at 540 nm.

### Cell culture

The CCD-19Lu normal human lung fibroblast cell line was purchased from the American Type Culture Collection (ATCC) and cultured in complete Eagle’s Minimum Essential Medium (LGC Standards, Teddington, Middlesex, UK) supplemented with 10 % heat-inactivated fetal calf serum (Serana, Pessin, Germany) and 1 % penicillin/streptomycin (Thermo Fisher Scientific). Primary human pulmonary fibroblasts (HPF) were purchased from PromoCell (Heidelberg, Germany) and cultured in complete Fibroblast Growth Medium 2 supplemented with 2.5 % fetal calf serum, 1 ng/mL recombinant human fibroblast growth factor, 5 µg/mL recombinant human insulin (PromoCell) and 1 % penicillin/streptomycin (Thermo Fisher Scientific). Cells were maintained at 37 °C in a humidified atmosphere containing 5 % CO_2_.

### TGF-β1- induced differentiation

Fibroblasts were seeded in 6-well plates and cultured for 24 h until circa 70 % confluence was reached. Cells were then serum-starved for 24 h and subsequently treated with recombinant TGF-β1 (5 ng/mL) for 72 h. Experiments were conducted under serum-free and antibiotic-free conditions. Controls were processed identically without TGF-β1.

### Reverse transcription and real-time PCR

Total RNA was extracted using the GeneJET RNA Purification Kit (Thermo Fisher Scientific) according to the manufacturer’s instructions. Then, total RNA (0.2 µg) was reverse transcribed using the RevertAid Moloney murine leukemia virus reverse transcriptase (Thermo Fisher Scientific). Quantitative real-time PCR was performed with the LightCycler 480 system (Roche Diagnostics, Mannheim, Germany), using PowerUp SYBR Green fluorescent mix (Thermo Fisher Scientific). Primer sequences (sense and antisense) are reported in Suppl. Fig. 1a. Relative gene expression was calculated using the ΔΔCt method with human ribosomal protein S16 (RPS16) as the reference gene.

### siRNA transfection

Specific predesigned siRNAs targeting LGMN, CatV, CysM/E and hCC, as well as the scrambled siRNA used as a control, were obtained from Qiagen (Hilden, Germany) (Suppl. Fig. 1b). After 72 h of TGF-β1 treatment, myofibroblasts were transfected with a 50 nM siRNA mixture using HiPerFect transfection reagent (Qiagen) in basal medium. The transfection mixture was applied to cells seeded in 6-well plates and replaced after 6 h. Treated cells were incubated in serum-free culture medium for 48 h prior to mRNA collection and for 72 h before protein extraction. All experiments were performed in triplicate.

### Pharmacological inhibitors

#### LGMN inhibitor (RR-11a)

myofibroblasts were incubated with 1 µM RR-11a in serum-free culture medium for 48 to 72 h following 72 h of TGF-β1 differentiation. Control experiments were performed in the absence of RR-11a. Inhibition of Smad-3 phosphorylation (SIS3): fibroblasts were seeded into 6-well plates and cultured for 24 h, followed by serum starvation for an additional 24 h. Subsequently, SIS3 (4 µM) was added for 1 h followed by TGF-β1 stimulation (5 ng/mL during 72 h). Control experiments were performed in the absence of SIS3 and in the absence of both SIS3 and TGF-β1.

### Wound healing assay

Fibroblasts were seeded into 24-well plates and differentiated with TGF-β1 as described above. Cell monolayers were scratched using a sterile pipette tip, washed with PBS and treated with or without RR-11a (1 µM) for 72 h. Wound closure was monitored every 6 h using the Incucyte SX5 Live-Cell Analysis System (Sartorius AG, Sartorius France, Dourdan, France). The area of the scratch was measured over time using ImageJ software and the relative closure was calculated based on the basal area (T0) of each wound.

### Cell viability and real-time cell death

#### MTS viability assay

Fibroblasts were seeded into 96-well plates and differentiated with TGF-β1 as described above and then incubated with RR-11a (0.5 – 5 µM) in serum-free culture medium for 48 or 72 h. Cell viability was assessed using CellTiter 96® AQueous One Solution Cell Proliferation Assay (MTS, Promega, Charbonniere-Les-Bains, France) following manufacturer’s instructions. Cell supernatants were replaced with 100 µL of serum-free culture medium containing 20 µL/well MTS reagent and plates were incubated at 37°C for 2 h. The absorbance was measured at 490 nm and results were normalized using myofibroblasts cultivated in absence of RR-11a as reference (100% of viability).

#### Cell death assay

Fibroblasts were seeded into 48-well plates and differentiated with TGF-β1 as described above. Cells were pretreated with RR-11a (1 µM) for 1 h before the addition of the apoptosis inducers: BV6 (20 µM) or staurosporine (STS, 0.5 µM) in serum-free culture medium containing 1 µM of PI. Cells were incubated in the IncuCyte SX5 live-cell analysis system and pictures were taken every hour with a red filter and bright-field. Positive, red-stained cells due to PI DNA-binding fluorescent indicating death were analyzed using the IncuCyte software package and results were expressed as the PI-positive area per image.

### Statistical analysis

Data are expressed as the median ± quartile range and analyzed statistically with the nonparametric Mann-Whitney test or the one-sample Wilcoxon test as mentioned (GraphPad Prism software version 9.5, GraphPad Software Inc., San Diego, CA, USA). Differences were considered statistically significant at p-value < 0.05 (*, p < 0.05; **, p < 0.01; ***, p < 0.001; ****, p < 0.0001).

## RESULTS

### LGMN, CatV and CysM/E are upraised in IPF lung tissue and BALF

To better refine the proteolytic landscape associated with pulmonary fibrosis, we first examined the expression of LGMN, CatV and CysM/E in lung biopsy samples obtained from non-IPF (n = 27) and IPF patients (n = 26), processed as described in the Experimental Section. RT-qPCR analysis confirmed the expected transcriptional upregulation of canonical fibrotic markers (α-SMA, collagen I), as well as fibronectin and elastin in IPF samples Suppl. Fig. 2a). Consistently, a noteworthy increase in their protein levels, apart from fibronectin whose expression level was only slightly enlarged, is found in IPF tissue lysates (Suppl. Fig. 2b-2g). ELISA and immunoblotting revealed a robust increase in immunoreactive LGMN in pooled IPF samples (Fig. 1a-1b). Densitometric analysis of individual samples indicated that the rise was significant for mature LGMN, whereas pro-LGMN levels remained unchanged (Suppl. Fig. 3a). Accordingly, LGMN activity, assessed using Ac-dTyr-Tic-Ser-Asn-ACC, a selective fluorogenic substrate, was significantly higher in IPF tissue lysates and was fully inhibited by MP-L01 [29], but not by E-64, the broad-spectrum inhibitor of cysteine cathepsins, thereby confirming the specificity of the enzymatic assay (Fig. 1c). Also, an overall increase of immunoreactive CatV and its proform was detected in IPF samples, as confirmed by ELISA (Fig. 1d-1e). WB analysis of individual samples supported that the increase in glycosylated mature CatV and pro-CatV, but not in their non-glycosylated forms [30], was significant in IPF samples (Suppl. Fig. 3b1-3b2). However, due to the current lack of a selective CatV substrate, its enzymatic activity could not be directly quantified. Likewise, CysM/E expression was increased in IPF tissue lysates, as observed for hCC (control) (Fig. 1f-1h). Interestingly, the level of 12 kDa non-glycosylated CysM/E form was significantly increased, whereas no significant change was observed for the 17 kDa form (Suppl. Fig. 3c). Enlarged levels of immunoreactive LGMN and CatV were detected in IPF BALF samples (Fig. 1i-1j). Likewise, we observed an increase of immunoreactive CysM/E in IPF BALFs (Fig. 1k-1l), as previously observed for hCC [18]. Nevertheless, the concentration of CysM/E (median value: ∼2.83 pg/µg of total protein) is approximately 20-fold lower than that of hCC (median value: ∼63.55 pg/µg of total protein) in the BALFs of IPF patients, suggesting a slighter contribution of extracellular CysM/E to the homeostasis of the cathepsin-dependent proteolytic balance.

**Fig. 1.**
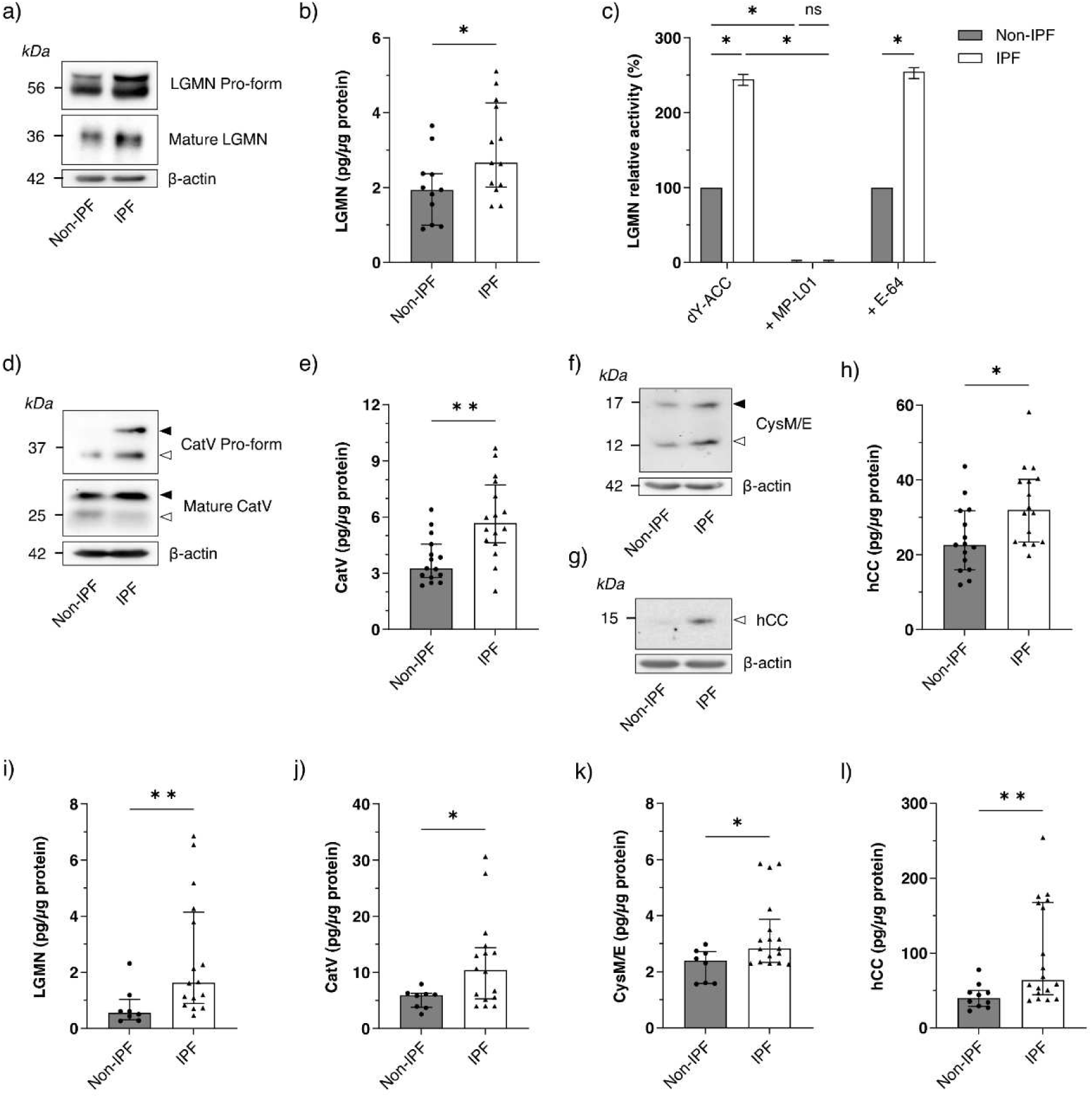
Expression levels of LGMN, CatV, CysM/E and hCC in human IPF *vs* non-IPF samples. (**a**) Western blot analysis of LGMN in pooled lung biopsy extracts from non-IPF or IPF patients (100 µg protein, β-actin as a loading control). (**b**) ELISA quantification of total human LGMN in individual lung biopsy extracts (non-IPF *vs* IPF). (**c**) LGMN peptidase activity measured using the dY-ACC fluorogenic substrate, with MP-L01 or E-64 (10 µM) used as inhibitors. The experiment was performed three times, each in duplicate. (**d**) Western blot analysis of CatV in pooled lung biopsy extracts (100 µg protein, β-actin as a loading control). Black arrows indicate glycosylated forms, whereas white arrows indicate non-glycosylated forms. (**e**) ELISA quantification of total human CatV in individual lung biopsy extracts. Western blot analysis of (**f**) CysM/E and (**g**) hCC in pooled lung biopsy extracts (100 µg protein, β-actin as a loading control). (**h**) ELISA quantification of total hCC in individual lung biopsy extracts. ELISA dosage of (**i**) LGMN, (**U**) CatV, (**k**) CysM/E and (**l**) hCC in individual human BALFs samples. All Western blot figures show a representative result from three independent experiments. Bars represent median ± quartile. Statistical significance was assessed using the Mann-Whitney test (* p<0.05; ** p<0.01)

### TGF-β1 regulates LGMN, CatV, and CysM/E expression in human lung fibroblasts

Given the scarcity of clinical specimens, we previously established a cellular model (primary human lung CCD-19Lu fibroblasts) for mechanistic studies of TGF-β1-driven myodifferentiation and related regulation of cysteine cathepsins [13, 20]. TGF-β1 treatment induced differentiation of CCD-19Lu fibroblasts (used as reference cells), as shown by enhancement of representative fibrotic markers (i.e., α-SMA and collagen I) at both the mRNA and protein levels (Suppl. Fig. 4a-4c). To strengthen these observations, we extended our study to a second cellular model (i.e., primary human pulmonary fibroblasts: HPF cells). For clarity and readability of the body text, data related to TGF-β1-treated HPF cells were reported as supplementary data (see Suppl. Fig. 5).

#### LGMN regulation

LGMN was transcriptionally (Fig. 2a) and translationally upregulated following TGF-β1 stimulation (Fig. 2b-2c). Likewise, the enzymatic activity of intracellular mature LGMN was significantly increased (Fig. 2d), as previously observed in IPF cell lysates (Fig. 1c). Also, TGF-β1 elicited secretion of pro-LGMN (56 kDa), while extracellular mature LGMN (36 kDa) was not detected (Fig. 2e-2f). Consistent with its striking stability at neutral pH, pro-LGMN may constitute a latent reservoir for generation of active LGMN under appropriate microenvironmental conditions [31]. In an analogous way, secreted CatB and CatL were mostly found as zymogens in lung fibroblasts [13].

**Fig. 2.**
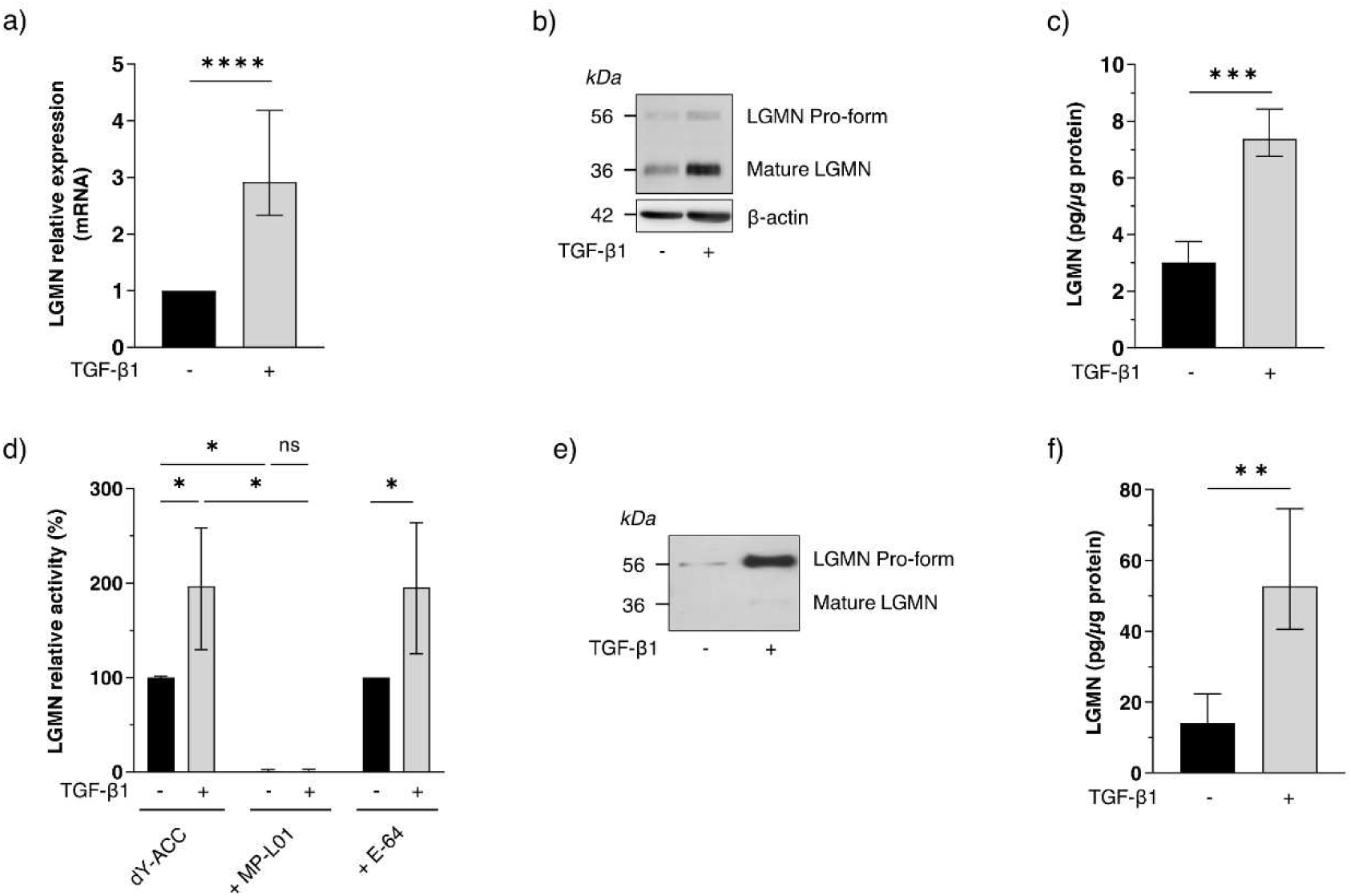
TGF-β1-induced myofibroblast differentiation increases LGMN expression and secretion in human pulmonary fibroblasts. CCD-19Lu cells were treated with recombinant human TGF-β1 (5 ng/mL) for 3 days to induce differentiation into myofibroblasts. (**a**) LGMN mRNA transcription was analyzed by RT-qPCR and expressed as the fold change in myofibroblasts relative to fibroblasts. LGMN protein levels in cell lysates were determined by (**b**) Western blot analysis (30 µg protein, β-actin as a loading control) and (**c**) ELISA quantification (n=6). (**d**) LGMN peptidase activity was measured in cell lysates (20 µg protein) using the dY-ACC fluorogenic substrate with MP-L01 or E-64 (10 µM) used as inhibitors. Experiments were performed in duplicate (n=4). Secreted LGMN in culture supernatant was analyzed by (**e**) Western blot (10 µg protein) and (**f**) ELISA quantification (n=6). A representative blot from three independent experiments is shown. Bars represent median ± quartile. Statistical significance was assessed using the Mann-Whitney test (* p<0.05; ** p<0.01; *** p<0.001), except for RT-qPCR data, which were analyzed using the one-sample Wilcoxon test (**** p<0.0001)

#### CatV regulation

Contrariwise, CatV mRNA and its intracellular protein level were reduced upon TGF-β1 stimulation (Fig. 3a-3b), differing from CatL and CatK, which are not transcriptionally regulated during TGF-β1-dependent myodifferentiation [14]. Accordingly, elastinolytic activity was decreased (Fig. 3c). The DQ-elastin hydrolysis was inhibited by E-64, but not by LHVS, a selective CatS pharmacological inhibitor (data not shown). As both pro-CatS and mature CatS were below the immunodetection threshold in CCD-19Lu myofibroblasts [14], this rules out significant contribution of CatS to fibroblast-associated elastinolysis. Likewise, present findings support that CatV is a major contributor to fibroblast-associated elastin degradation under these experimental conditions.

**Fig. 3.**
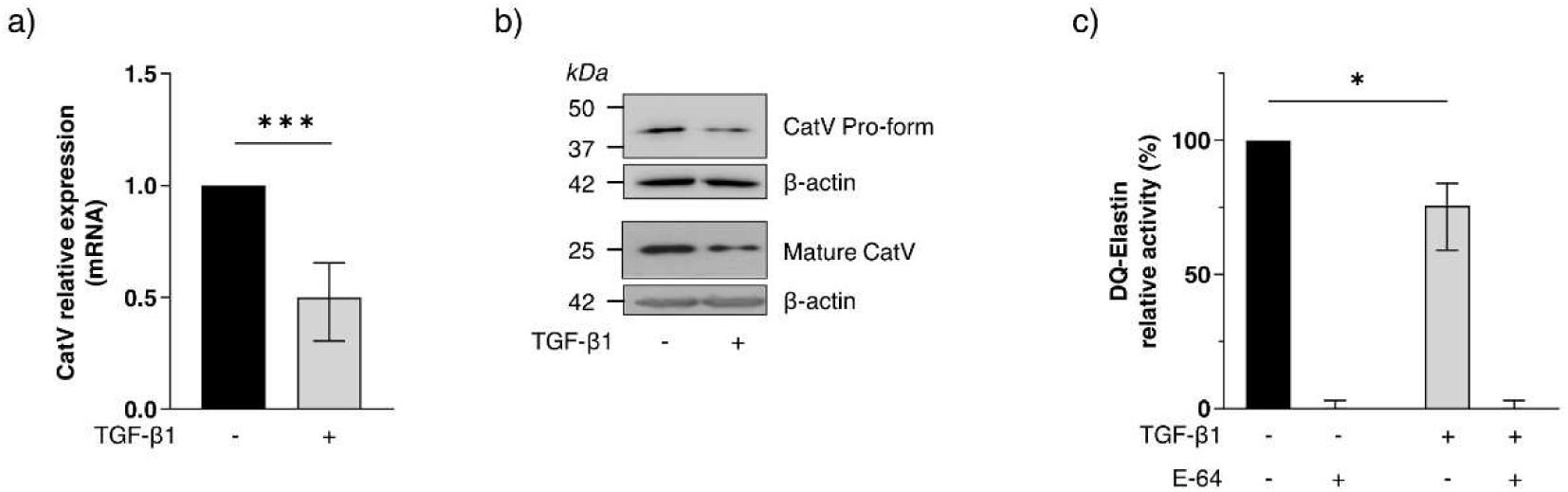
TGF-β1-induced myofibroblast differentiation decreases CatV expression and elastinolytic activity in human pulmonary fibroblasts. CCD-19Lu cells were treated with recombinant human TGF-β1 (5 ng/mL) for 3 days to induce differentiation into myofibroblasts. (**a**) CatV mRNA transcription was analyzed by RT-qPCR (n=6) and expressed as the fold change in myofibroblasts relative to fibroblasts. (**b**) CatV protein level in cell lysates was analyzed by Western blot (50 µg protein, β-actin as a loading control). All Western blot figures show a representative result from three independent experiments. (**c**) Elastinolytic activity was measured in cell lysates (100 µg protein) using the DQ-Elastin fluorogenic substrate. Assays were performed in the presence of LHVS (10 nM), with or without E-64 (10 µM), (n=4). Bars represent median ± quartile. Statistical significance was assessed using the one-sample Wilcoxon test (* p<0.05; *** p<0.001)

#### CysM/E regulation

TGF-β1 makedly upregulated CysM/E transcription (Fig. 4a), conversely to both hCC and stefin B (type 1 cystatin) whose mRNA levels remained unchanged, as demonstrated earlier [13]. Regarding protein expression, TGF-β1 also increased the secretion of both CysM/E and hCC (Fig. 4b-4e), hence enhancing the inhibitory potential of fibroblast secretome toward extracellular cysteine cathepsins [13]. However, the concentration of secreted CysM/E (median value: ~4.8 pg/µg of total protein) is circa 60-fold lower than that of hCC (median value: ~290.5 pg/µg of total protein), consistent with the relative abundance observed in BALF samples from IPF patients. Similar results were obtained in HPF cells with respect to the transcription, intracellular protein expression and secretion of LGMN, CatV, CysM/E and hCC as well as LGMN activity, underlining the relevance of fibroblast differentiation models (Suppl. Fig. 5d-5m).

**Fig. 4.**
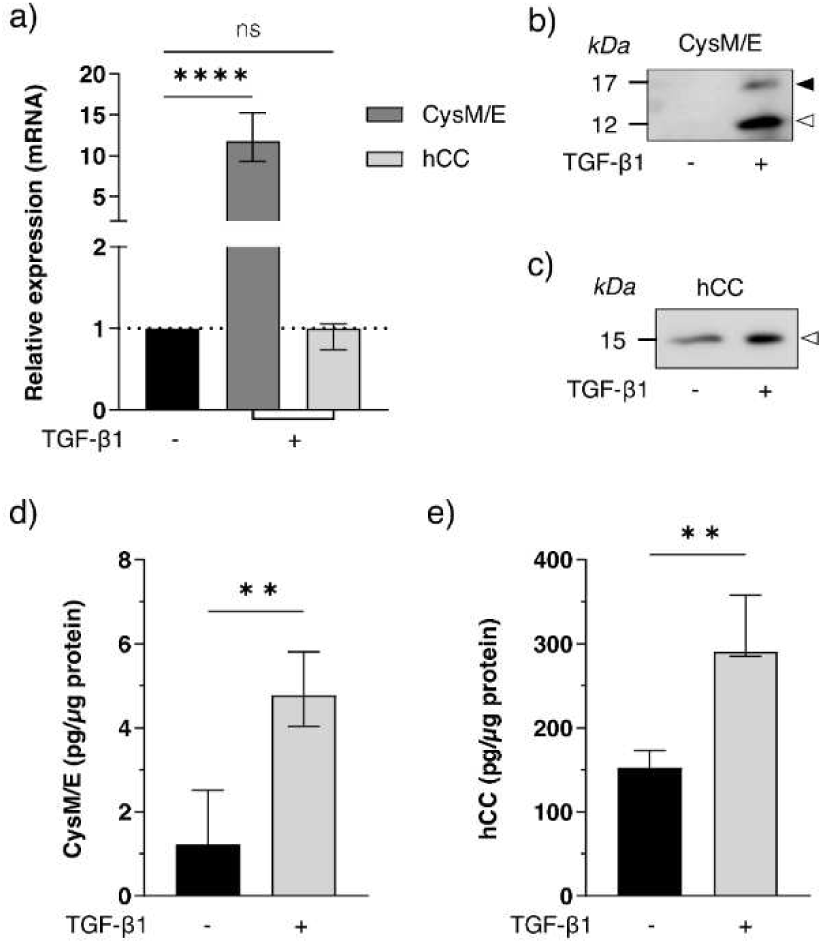
TGF-β1-induced myofibroblast differentiation regulates cystatin secretion by human pulmonary fibroblasts. CCD-19Lu cells were treated with recombinant human TGF-β1 (5 ng/mL) for 3 days to induce differentiation into myofibroblasts. (**a**) CysM/E and hCC mRNA expressions were analyzed by RT-qPCR (n=6) and expressed as the fold change in myofibroblasts relative to fibroblasts. Western blot analysis of secreted (**b**) CysM/E (10 µg protein) and (**c**) hCC (5 µg protein) in culture supernatant. Black arrows indicate glycosylated forms, whereas white arrows indicate non-glycosylated forms. All Western blot figures show a representative result from three independent experiments. ELISA quantification of secreted (**d**) CysM/E (glycosylated and non-glycosylated forms) and (**e**) hCC in culture supernatant (n=5). Bars represent median ± quartile. Statistical significance was assessed using the one-sample Wilcoxon test for RT-qPCR (ns: not significant; *** p<0.001) and the Mann-Whitney test for ELISA quantification (** p<0.01)

### Effect of LGMN, CatV, CysM/E and hCC silencing on fibroblast phenotype

CCD-19Lu fibroblasts were transfected with specific siRNAs targeting hCC, LGMN, CatV, or CysM/E. Results of hCC silencing (transcriptional and translational expression levels) were reported earlier [13]. Effective knockdown of LGMN, CatV and CysM/E was confirmed at both mRNA and protein levels (Suppl. Fig. 6a-6c). Otherwise, our transfection attempts were unsuccessful for HPF cells (data not shown). Next, we evaluated the respective impact of LGMN, CatV, or CysM/E silencing on fibroblast differentiation by assessing the expression level of α-SMA and profibrotic hCC [17] (Suppl. Fig. 7). Genetic impairment of LGMN and CatV did not affect α-SMA protein expression, endorsing that LGMN and CatV are not involved in TGF-β1-induced myofibroblast differentiation, conversely to that observed following siRNA-mediated down-regulation of CatB [13]. Similarly, CysM/E silencing did not alter α-SMA expression (Suppl. Fig. 7a) suggesting that this inhibitor is not directly involved in the regulation of the myofibroblastic phenotype. Also, hCC secretion was not altered by silencing of LGMN, CatV, or CysM/E (Suppl. Fig. 7b-7c). Because LGMN deficiency has been proposed to affect maturation of CatB and CatL from single-chain forms into their two-chain forms in murine models [32], we examined CatB and CatL processing in LGMN-silenced human lung fibroblasts (Fig. 5a-5f). Neither intracellular and secreted CatB and CatL protein levels, nor both CatB- and CatL-related peptidase activities were altered by LGMN knockdown, advocating that regulatory differences could depend on cell type and/or species specificity.

**Fig. 5.**
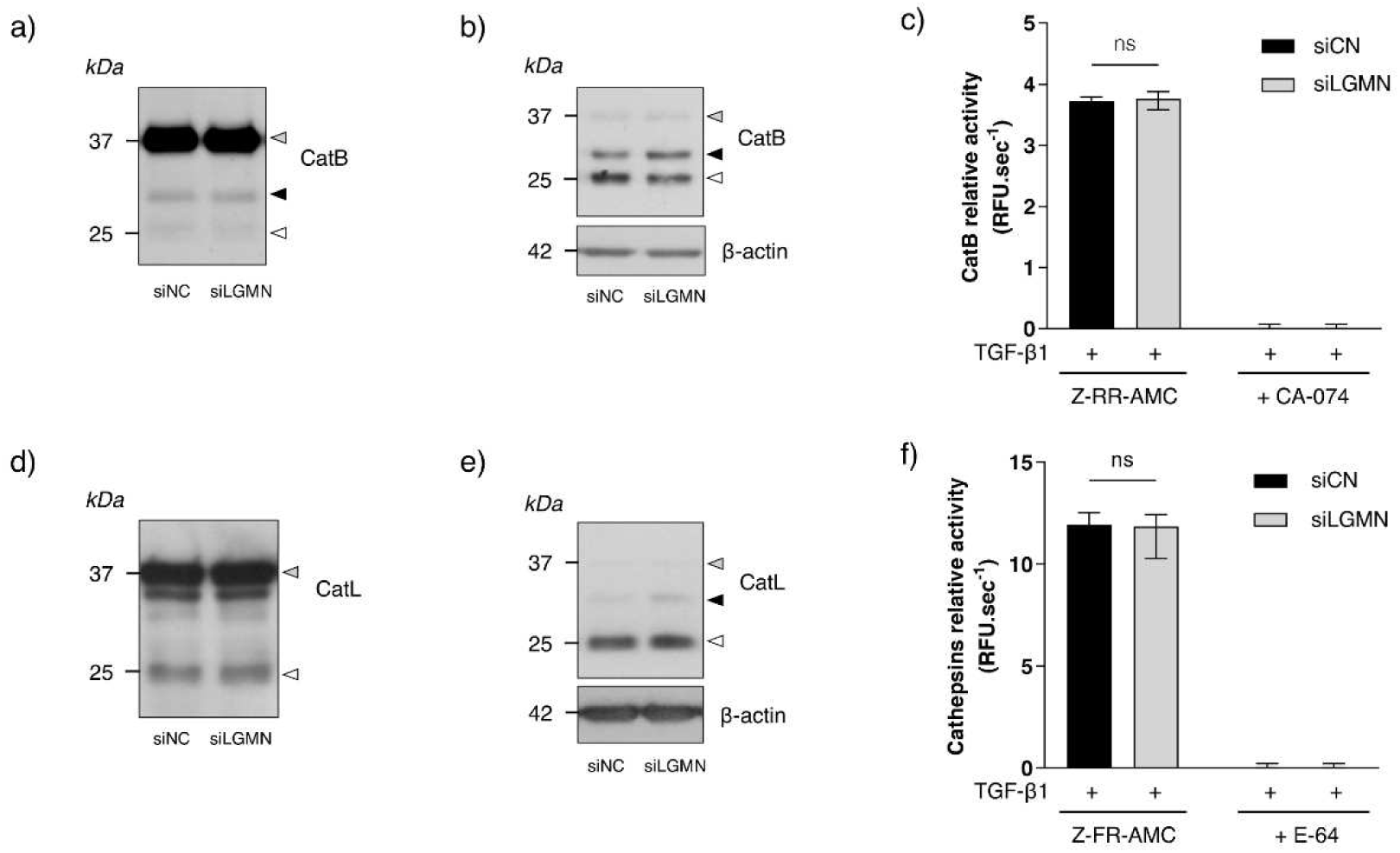
Effect of LGMN silencing on CatB and CatL maturation in human pulmonary myofibroblasts. CCD-19Lu cells were differentiated into myofibroblasts with recombinant human TGF-β1 (5 ng/mL) for 3 days. Cells were then transfected with a siRNA targeting LGMN (siLGMN) or a scrambled siRNA used as a negative control (siNC). Western blot analysis of (**a**) secreted CatB (10 µg protein) and (**b**) intracellular CatB (20 µg protein, β-actin as a loading control). Grey arrows indicate the zymogen, black arrows indicate the single-chain forms and white arrows indicate the two-chain forms. (**c**) CatB peptidase activity was measured in cell lysates (10 µg protein) using the Z-RR-AMC fluorogenic substrate, with CA-074 (10 µM) used as a selective inhibitor. Experiments were performed in duplicate (n=4). Western blot analysis of (**d**) secreted CatL (10 µg protein) and (**e**) intracellular CatL (20 µg protein, β-actin as a loading control). (**f**) Overall cathepsins B/L-like peptidase activity was measured in cell lysates (5 µg protein) using the Z-FR-AMC fluorogenic substrate, with E-64 (10 µM) used as an inhibitor. Experiments were performed in duplicate (n=4). All Western blot figures show a representative result from three independent experiments. Bars represent median ± quartile. Statistical significance was assessed using the Mann-Whitney test (ns: not significant)

### LGMN and CatV contribute to fibronectin and elastin remodeling

TGF-β1 significantly increased fibronectin and elastin mRNA and protein expression levels (Fig. 6a-6c), as reported for conventional fibrotic markers (i.e., collagen and α-SMA). To determine whether CatV and LGMN modulate ECM turnover, we assessed the effect of protease silencing on matrix accumulation. As anticipated, silencing of either CatV or LGMN did not alter collagen levels (data not shown), corroborating that both CatV and LGMN did not act as collagenases. Accordingly, silencing of CysM/E, the dual inhibitor of CatV and LGMN, did not decrease accumulation of type I collagen (data not shown), in contrast to the knockdown of hCC, the primary extracellular inhibitor of cysteine cathepsins, which led to a pronounced reduction of soluble and insoluble collagens levels [20]. Then, we scrutinized the ability of LGMN and CatV to modulate the extracellular accumulation of fibronectin and elastin, as well as the likely impact of both hCC and CysM/E.

**Fig. 6.**
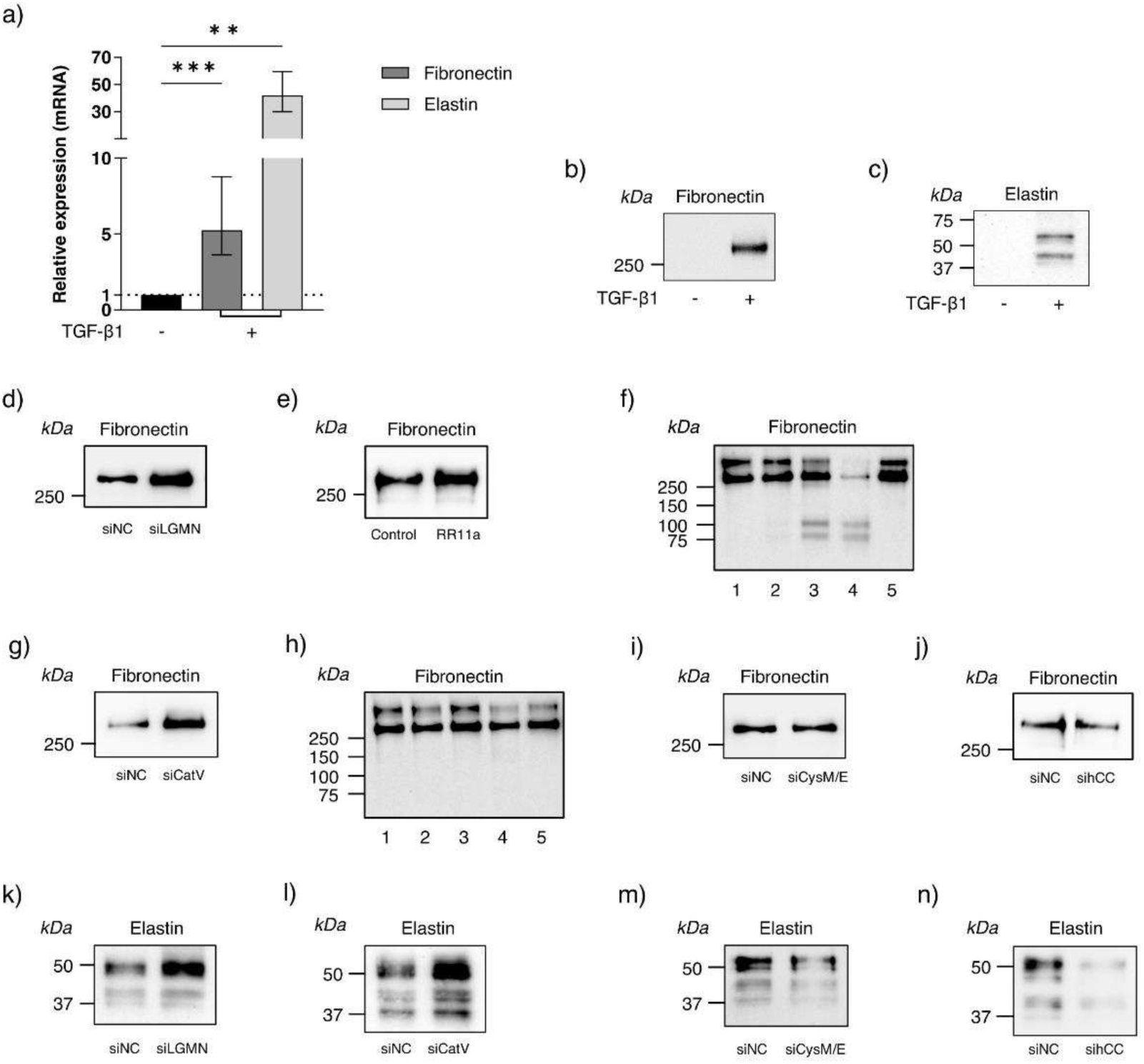
Effects of LGMN, CatV, CysM/E and hCC silencing or inhibition on extracellular fibronectin and elastin levels. CCD-19Lu cells were treated with recombinant human TGF-β1 (5 ng/mL) for 3 days to induce differentiation into myofibroblasts. (**a**) Fibronectin and elastin mRNA expression was analyzed by RT-qPCR (n=6) and expressed as the fold change in myofibroblasts relative to fibroblasts. Western blot analysis of (**b**) fibronectin and (**c**) elastin in culture supernatant (10 µg protein). On day 3, myofibroblasts were transfected with a siRNA targeting LGMN, CatV, CysM/E or hCC or with a scrambled siRNA used as negative control (siNC). Western blot analysis of fibronectin following (**d**) LGMN silencing or (**e**) treatment with the LGMN inhibitor RR-11a (1 µM). (**f**) *in vitro* hydrolysis of human recombinant fibronectin by LGMN was analyzed by immunoblotting (lane 1: fibronectin control without enzyme; lanes 2 – 4: enzyme to substrate molar ratio 1:10,000; 1:1000; 1:100 respectively; lane 5: molar ratio 1:100 in the presence of RR-11a (10 µM)). (**g**) Western blot analysis of fibronectin by myofibroblasts following CatV silencing. (**h**) *in vitro* hydrolysis of human recombinant fibronectin by CatV was analyzed under the same experimental conditions as in (f), except that E-64 (10 µM) was used as the inhibitor in lane 5. Western blot analysis of fibronectin by myofibroblasts following (**i**) CysM/E silencing or (**U**) hCC silencing. Western blot analysis of elastin by myofibroblasts following (**k**) LGMN, (**l**) CatV, (**m**) CysM/E, or (**n**) hCC silencing. All Western blot figures show a representative result from three independent experiments. Bars represent median ± quartile. Statistical significance was assessed using the one-sample Wilcoxon test (** p<0.01; *** p<0.001)

#### Fibronectin

LGMN silencing, as well as pharmacological inhibition with RR-11a, increased fibronectin deposition (Fig. 6d-6e), while LGMN cleaved recombinant fibronectin *in vitro* (Fig. 6f), which confirms that LGMN functions as a fibronectin-degrading enzyme, as suggested in a murine model [33, 34]. CatV silencing also enhanced fibronectin accumulation (Fig. 6g); however, recombinant CatV did not cleave fibronectin *in vitro* (Fig. 6h), suggesting an indirect proteolytic pathway, possibly through activation of downstream proteolytic pathways or modulation of ECM-associated proteases. Genetic impairment of hCC promoted hydrolysis of fibronectin, whereas CysM/E silencing had no substantial effect (Fig. 6i-6j).

#### Elastin

Deficiency of LGMN or CatV led to elastin accumulation (Fig. 6k-6l). Conversely, silencing of hCC - and to a lesser degree CysM/E - reduced elastin levels (Fig. 6m-6n), strengthening that both proteases, especially CatV, contribute to elastin turnover during fibroblast differentiation.

### Smad-3-driven regulation of LGMN and CysM/E

Given that TGF-β1 upregulates LGMN and CysM/E transcription and downregulates CatV transcription, we investigated whether this mechanism could involve the Smad-3 signaling pathway, a main downstream effector of TGF-β1 during FMT. Accordingly, fibroblasts were treated with SIS3, a potent and specific inhibitor of Smad-3 phosphorylation with antifibrotic properties [35]. SIS3 did not impact CatV mRNA expression; in contrast, LGMN transcription and enzymatic activity were reduced in the presence of SIS3 (Fig. 7a-7c), advocating that LGMN transcription is regulated upstream by Smad-3 signaling. As anticipated, SIS3 had no effect on hCC mRNA level (Fig. 7d), which corroborates that hCC transcription does not depend on the Smad-3 pathway [13]. Contrariwise, CysM/E transcription was markedly decreased (Fig. 7e). Consequently, the peptidase activity of secreted cysteine cathepsins increased in conditioned media from SIS3-treated cells (i.e., corresponding to fibroblast secretome), consistent with reduced CysM/E expression and decreased extracellular inhibition of cysteine proteases (Fig. 7f). Taken together, these results established that TGF-β1 drives LGMN and CysM/E expression, via the Smad3 signaling pathway. Conversely, Smad-3 does not trigger upstream CatV transcription during TGF-β1-dependent differentiation of lung fibroblasts, as established for CatL and CatK [14].

**Fig. 7.**
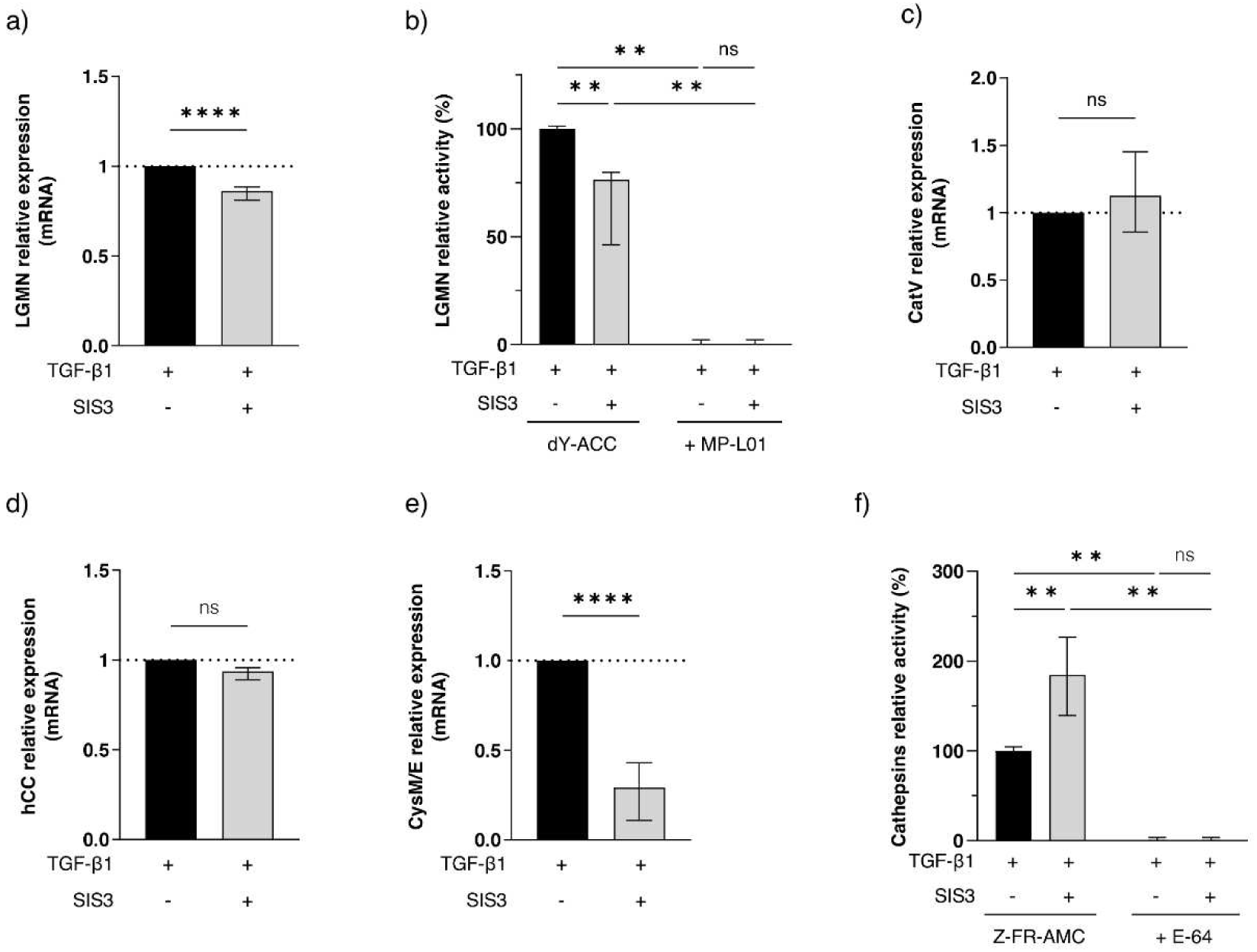
Effect of Smad-3 inhibition during TGF-β1-induced myofibroblast differentiation on LGMN, CatV and cystatins expression. CCD-19Lu cells were treated with recombinant human TGF-β1 (5 ng/mL) for 3 days in presence or absence of SIS3 (4 µM), an inhibitor of Smad-3 phosphorylation. (**a**) LGMN mRNA expression was analyzed by RT-qPCR (n=4). Data are expressed as fold change in treated myofibroblasts relative to control cells. (**b**) LGMN peptidase activity in cell lysates (20 µg protein) was measured using the dY-ACC fluorogenic substrate, with MP-L01 (10 µM) used as inhibitor. Experiments were performed in duplicate (n=4). (**c**) CatV, (**d**) hCC, and (**e**) CysM/E mRNA expressions were analyzed by RT-qPCR. (**f**) Overall cathepsins B/L-like peptidase activity in culture supernatant (5 µg protein) was measured using the Z-FR-AMC fluorogenic substrate, with E-64 (10 µM) used as inhibitor. Experiments were performed in duplicate (n=4). Bars represent median ± quartile. Statistical significance was assessed using the one-sample Wilcoxon test (ns: not significant; ** p<0.01; **** p<0.0001) for RT-qPCR and the Mann-Whitney test (** p<0.01) for enzymatic activity

### LGMN promotes fibroblast-mediated wound closure

Given its role in fibronectin remodeling, we examined whether LGMN could be involved in ECM dynamics and contributes functionally to fibroblast migration. Although most mechanistic experiments were performed using CCD-19Lu fibroblasts, scratch wound-healing assays were conducted in HPF cells, because CCD-19Lu monolayers poorly tolerated mechanical injury, exhibited limited post-scratch expansion, and failed to reproducibly recover after wounding. Importantly, as shown above, both cellular models displayed similar transcriptional and translational responses to TGF-β1 stimulation, supporting the relevance of HPF cells for functional migration analyses. We first performed a MTS assay on cells treated with RR-11a, a LGMN synthetic inhibitor, to evaluate its potential cytotoxicity and impact on cell viability. No deleterious effects were observed (Suppl. Fig. 8a). In addition, HPF cells were treated with BV6, a pharmacological mimetic of SMAC (Second Mitochondria-derived Activator of Caspases) and an antagonist of cIAP1 (cellular Inhibitor of Apoptosis Protein-1) and XIAP (X-Linked Inhibitor of Apoptosis Protein) (Suppl. Fig. 8b-8c) or with staurosporine, a pan-inhibitor of protein kinases and an activator of the intrinsic apoptosis pathway (Suppl. Fig. 8d-8e). Analysis of propidium iodide fluorescence demonstrated that RR-11a did not induce significant increase in cell death. Scratch wound-healing assays revealed that pharmacological inhibition of LGMN with RR-11a significantly delayed wound closure during the first 48 h post-injury (Fig. 8a-8b). Wound closure ultimately normalized after ~54 h, indicating a transient but functionally meaningful impairment of myofibroblasts migration. Because efficient wound healing depends on continuous ECM remodeling and a dynamic microenvironment, these findings support an operational role for LGMN in ECM reorganization, likely through the proteolytic remodeling of fibronectin required for efficient myofibroblast motility and migration.

**Fig. 8.**
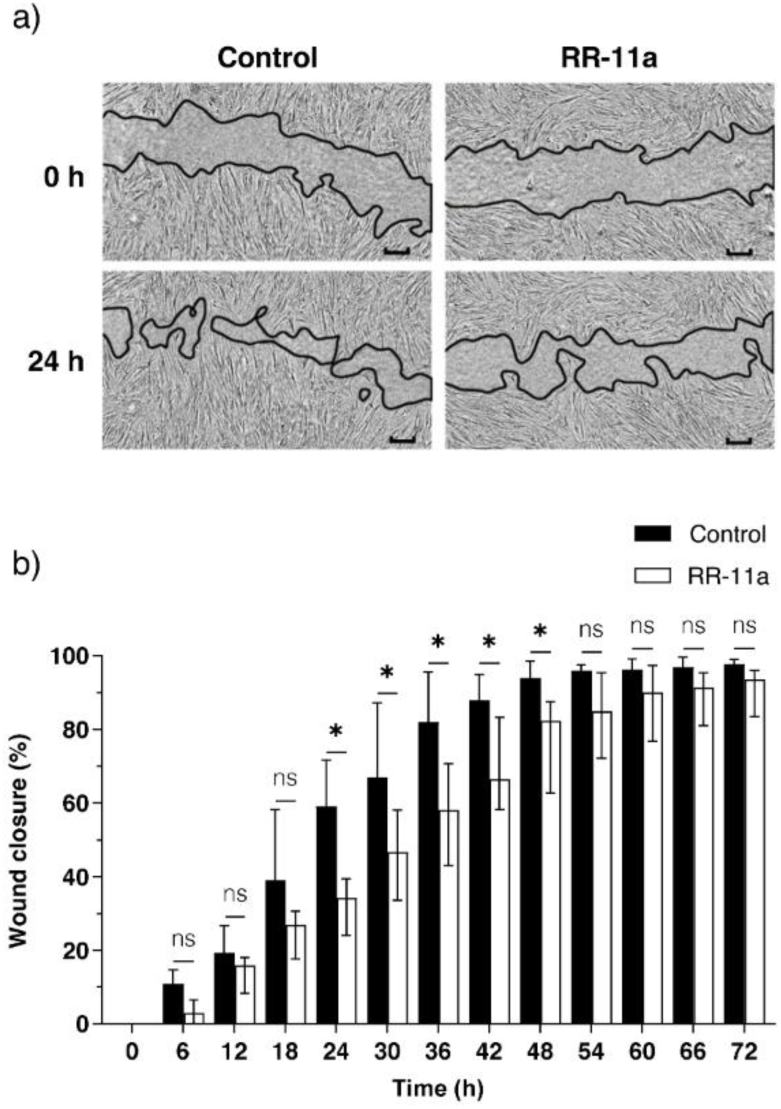
Effect of LGMN inhibition on wound closure capacity of pulmonary myofibroblasts. TGF-β1-differentiated HPF cells were subjected to a scratch assay and wound healing process was monitored over 72 h in presence or absence of RR-11a (1 µM). (**a**) Representative images of wound healing assay at 0 and 24 h (scale bar = 500 µm). (**b**) Percentage of wound closure over time relative to the wound area at T0. Experiments were performed in triplicate (n=3). Bars represent the median ± quartile. Statistical significance was assessed using the Mann-Whitney test (ns: not significant; * p<0.05)

## DISCUSSION

Pulmonary fibrosis is driven by excessive ECM deposition and profound remodeling of lung architecture, processes tightly regulated by a network of proteases and antiproteases [6]. While metalloproteinases and serine proteases have well-established roles in this context [4, 5, 36], cysteine proteases - particularly cathepsins B, L, K and S - and their endogenous inhibitors have emerged as key contributors to fibrotic progression [14, 17, 37, 38]. In this study, we focused on the less-explored cysteine proteases LGMN and CatV, together with their dual inhibitor CysM/E, to clarify their functional roles in ECM turnover during pulmonary fibrosis.

We first established the expression profiles of LGMN, CatV, and CysM/E. We demonstrated that LGMN, CatV, and CysM/E are significantly increased in IPF lung biopsies and BALFs *vs* non-IPF specimens. Notably, glycosylated forms of CatV were predominantly elevated, consistent with prior evidence showing that N-glycosylation promotes CatV subcellular localization, secretion, stability and pericellular proteolytic activity [39–41]. Glycosylated CatV was correlated with proteolytic activity and invasive phenotypes in lung tumors [42]. Moreover, glycosylated pro-CatV partly retains catalytic competence in human cells and tissues as shown by means of DCG-04, a broad-spectrum activity-based probe of cysteine cathepsins [43], suggesting that IPF lungs could accumulate a partially active pool of elastinolytic CatV capable of contributing to ECM remodeling. However, in the absence of a potent and selective CatV substrate, no direct CatV activity could be quantified or uncovered in the present study. In addition to hCC, a presumed fibrosis biomarker [17, 18], an increase of CysM/E was observed in both lung specimens and BALFs of IPF patients. However, the concentration of CysM/E is low compared to that of hCC, suggesting that CysM/E would have an auxiliary role, while hCC constitutes the predominant extracellular cysteine cathepsin inhibitor in fibrotic lungs. CysM/E was preferentially detected as its non-glycosylated form (12 kDa) in IPF lung samples, while the protein level of the glycosylated form (17 kDa) did not differ between IPF and non-IPF biopsies. Nevertheless, the functional consequences are doubtless limited, since both glycosylated and non-glycosylated CysM/E display a similar inhibitory potency [44, 45]. Probably more importantly, CysM/E also inhibits auto-activation of secreted pro-LGMN, which exerts a finely tuned regulatory role in controlling both intra-and extracellular LGMN trafficking, processing and activity [46, 47]. Consistently, extracellular pro-LGMN can be internalized and processed into active intracellular forms, whereas internalization of extracellular CysM/E may distinctly reduce intracellular LGMN activity and cell migration [47, 48]. Taken together, data support that IPF lungs harbor increased levels of both LGMN and CatV and their natural inhibitors (hCC and CysM/E), leading to an altered protease-inhibitor balance.

In a second step, we carried out their in-depth transcriptional and translational characterization, using human lung fibroblasts (CCD-19 Lu and HPF cells). We established that TGF-β1 induces a coordinated regulation of LGMN, CatV and CysM/E. Also, data endorse that these two models of fibroblast differentiation are relevant for deciphering the functional role of CatV, LGMN. Plasma LGMN level was confirmed as a putative IPF biomarker [25], while LGMN contributes to fibronectin degradation, cell migration as well as tissue repair in various cell and animal models (for review: [24, 33, 34]. However, its role in fibrosis appears to be critically dependent on the physiological context. LGMN deficiency increases the secretion of profibrotic cytokines, whereas supplementation by ectopic LGMN showed a therapeutical benefit [49]. While LGMN deficiency reduces the severity of pancreatic fibrosis, LGMN could act as a profibrotic factor by participating, via the processing of pro-MMP-2 to active MMP-2, in the maturation of latent TGF-β1 precursor [24]. Conversely, in a model of obstructive nephropathy, macrophage-derived LGMN displayed a potent antifibrotic role through promotion of ECM turnover, while its deficiency aggravated renal fibrosis [50]. Moreover, TGF-β1 promotes both internalization and activation of extracellular pro-LGMN to catalytically active LGMN during osteoblast maturation [51], suggesting the involvement of a peculiar “TGF-β1-LGMN axis”. Thus, current studies support that the contrasting role of LGMN is most probably tissue dependent. Here, we demonstrated that TGF-β1 triggers LGMN transcription and upregulates intracellular mature and enzymatically active LGMN, while also promoting secretion of pro-LGMN, which is unexpectedly stable at neutral pH and thus may serve as a latent reservoir of activatable LGMN.

CatV is a human cysteine protease with strong elastinolytic activity, which probably plays a distinct role in remodeling of the fibroblast microenvironment [22]. In cutaneous scleroderma, serum level of CatV is decreased, while TGF-β1 reduces CatV expression in dermal fibroblasts, favoring ECM deposits [52]. However, elastin degradation may also release elastokines that fuel pro-fibrotic loops [53]. Consequently, CatV can promote ECM clearance or be pro-fibrogenic, depending on the local protease/antiprotease balance, pericellular pH, and cell source [54]. Here, CatV transcription and intracellular protein levels as well as elastinolytic activity were decreased by TGF-β1, in accordance with observations that CatV repression correlates with enhanced matrix deposition, contributing to skin fibrosis [52]. Of note, the decreased TGF-β1-dependent elastinolytic activity did not relate to CatS, which is very slightly produced by lung fibroblasts [14]. Moreover, the apparent discrepancy of CatV expression level between primary lung fibroblasts and IPF lung biopsies probably relies on the presence in IPF specimens of multiple cell types - in addition to fibroblastic cells - as shown by spatial transcriptomics analyses, integrating IPF scRNA-seq atlas data [55].

Alongside the confirmation that TGF-β1 does not drive hCC transcription, but promotes its secretion by myofibroblasts [13], we established here that CysM/E was prominently induced by TGF-β1, at both mRNA and protein levels. Accordingly, CysM/E upregulation could enhance the inhibitory potential of the fibroblast secretome with respect to its target proteases. However, due to its low abundance compared to that of hCC, the inhibitory function of CysM/E is probably auxiliary, playing a supportive role for secreted hCC, which is the predominant broad-spectrum inhibitor of LGMN and cysteine cathepsins.

As the major transcriptional mediator of TGF-β1 signaling in fibroblasts, Smad-3 triggers numerous profibrotic genes once phosphorylated. Consequently, Smad-3 deficiency attenuates fibrosis and alters wound repair dynamics [56, 57]. Smad-3 inhibition by SIS3, a pharmacological inhibitor of Smad-3 phosphorylation markedly reduced LGMN and CysM/E transcription and decreased LGMN activity, establishing that both genes are under the control of the canonical TGF-β1/Smad-3 axis. Conversely, treatment of fibroblasts with SIS3 did not affect either CatV or hCC mRNA levels. Data confirmed that hCC transcription is not regulated upstream by Smad-3 [13]. Likewise, Smad-3 did not drive CatV (CatL2) transcription, as reported for CatL and CatK during lung myodifferentiation [14]. This result advocates that regulation of cathepsin L-related proteases most likely obeys a distinct regulatory program that differs from that of CatB, which is involved in the TGF-β1-dependent fibroblast differentiation [13]. Interestingly, a correlation between LGMN and Smad3 expression was reported in macrophages and foam cells within atherosclerotic plaques. Subsequently, authors have proposed that a functional LGMN/Smad3 regulatory axis could be involved in pathological tissue remodeling frameworks [58]. Hence, present findings position LGMN and CysM/E as Smad-3-dependent components of the fibrotic response, whereas CatV appears regulated through a Smad-3-independent mechanism in lung myofibroblasts.

We next assessed the effects of LGMN, CatV, CysM/E, and hCC silencing on fibronectin and elastin accumulation, both of which were increased following TGF-β1 treatment. Silencing or pharmacological inhibition of LGMN resulted in increased fibronectin accumulation. Otherwise, LGMN efficiently cleaved fibronectin *in vitro*, substantiating that it could be a fibronectin-degrading enzyme of biological relevance. Although CatV did not directly hydrolyze fibronectin under the same experimental conditions, CatV silencing in fibroblasts also increased fibronectin expression level, which sustains an indirect regulation of fibronectin turnover. Although the underlying proteolytic mechanism remains elusive at this stage, a recent study has revealed that an antibody targeting CatV impairs fibronectin cleavage [59]. Elastin homeostasis was similarly affected. Likewise, CatV silencing increased elastin accumulation, consistent with its reported elastinolytic activity. On the other hand, LGMN knockdown also elicited an increase of elastin which supports that LGMN, which is not an elastase, could be involved upstream in elastolysis. One possible path could occur via the primary activation of pro-MMP-2 to mature MMP-2 [24]. In accordance with this hypothesis, LGMN participated in cardiac ECM remodeling by triggering the MMP-2 pathway [34]. Present data reveal that LGMN and CatV participate in protease networks that regulate fibronectin and elastin turnover. Also, hCC silencing forcefully decreased elastin levels, sustaining a pivotal role for cysteine cathepsins, including CatV, in elastinolysis. CysM/E silencing induced a similar but milder effect, which strengthens its secondary role compared to prevalent hCC. Interestingly, neither LGMN nor CatV inhibition altered α-SMA expression, in contrast to CatB, a proteolytic mediator of TGF-β1-induced lung myofibrogenesis and myofibroblast differentiation [13]. Similarly, CysM/E silencing did not modulate α-SMA expression. Functionally, present data reveal that LGMN and CatV contribute to ECM remodeling but do not directly participate in fibroblast activation.

A key operative outcome of LGMN activity emerged from wound-healing assays. Pharmacological inhibition of LGMN significantly delayed fibroblast migration during the early phases of scratch closure. Because cell migration requires dynamic and spatially controlled ECM remodeling, these results provide functional support for a model in which LGMN promotes cell migration by modulating fibronectin turnover during the healing phase. The transient nature of the migratory defect is consistent with the existence of compensatory proteolytic mechanisms that gradually restore ECM remodeling capacity. Therefore, TGF-β1, relayed by Smad-3, could be a key upstream regulator of LGMN enzymatic activity in fibrotic deposition as well as in tissue repair.

Overall, our findings delineate a dual regulatory axis in which TGF-β1 activates Smad-3 to induce LGMN and CysM/E expression, while repressing CatV independently of Smad-3. In this framework, LGMN and CatV act as ECM-modifying enzymes whose proteolytic activities promote fibronectin and elastin degradation, thereby contributing to tissue remodeling. The combination of hCC and CysM/E, by inhibiting LGMN and CatV, participates in the counter-regulatory mechanisms that favor ECM accumulation in fibrosis. The elevation of LGMN and CatV in IPF tissues could likely reflect an attempted compensatory response to excessive matrix deposition, yet one that is hindered by hCC and to a lesser extent by CysM/E. We identify LGMN as a Smad-3-dependent effector of ECM remodeling and myofibroblast migration that does not directly promote fibrogenesis, in contrast to other proteases such as CatB which promotes fibroblast differentiation. Together with CatV, LGMN contributes to fibronectin and elastin turnover and supports fibroblast motility. Their dysregulation, combined with altered expression of cystatins, could shape the pathological ECM landscape characteristic of IPF. These findings highlight LGMN and CatV as major proteolytic partners of ECM remodeling and suggest that modulating their proteolytic balance could offer novel avenues for therapeutic intervention in pulmonary fibrosis.

## CONCLUSION

In this study, we identify LGMN and CatV as previously under-recognized regulators of ECM dynamics during pulmonary fibrosis. Expression levels of both proteases are increased in IPF tissues and BALFs but exhibit distinct regulatory patterns in lung fibroblasts. TGF-β1, acting through Smad-3, drives LGMN transcription, intracellular activation, and secretion of its pro-form, while simultaneously inducing the dual inhibitor cystatin M/E (CysM/E). In contrast, despite being present in fibrotic lungs, CatV is repressed by TGF-β1 through a Smad-3-independent mechanism. Functional analyses demonstrate that LGMN and CatV contribute primarily to ECM remodeling rather than to fibroblast-to-myofibroblast differentiation. Loss of either enzyme increases fibronectin and elastin accumulation, whereas their endogenous inhibitors - especially hCC - favor matrix deposition. LGMN directly cleaves fibronectin and supports fibroblast migration, consistent with a role in dynamic ECM turnover required for tissue repair. These findings identify a cystatin-regulated LGMN/CatV partnership, in which TGF-β1/Smad-3-dependent induction of LGMN and CysM/E, together with altered CatV expression, shifts the protease–inhibitor balance and collectively shapes the fibrotic microenvironment. LGMN and CatV therefore emerge as mechanistically relevant effectors of ECM remodeling and may constitute potential therapeutic targets for modulating matrix homeostasis in fibrotic lung disease.

## Acknowledgements

We thank Dr Benoît Briard (University of Tours & INSERM, UMR1100, Research Center for Respiratory Diseases, Tours, France) for his gift of BV6 and staurosporine. Morpholinourea-leucinyl-homophenylalanine-vinyl-sulfone phenyl (LHVS) was generously provided by Prof. James H. McKerrow (Skaggs School of Pharmacy and Pharmaceutical Sciences, University of California, San Diego, CA, USA). We acknowledge Prof. Marcin Drag (Centre for Chemical Biology, Institute of Physical Chemistry, Polish Academy of Sciences, Warszawa, Poland) for his kind gift of legumain substrate and inhibitor.

## Contributions

Conceptualization: G.L., A.S.; methodology: B.R., G.L., A.S.; formal analysis: B.R., A.D., F.V., S.M-A., G.L., A.S.; investigation: B.R., A.D., R.A., C.L., L.V.; resources: D.S., F.L., M.P., S.M-A.; data collection: D.S, S.M-A.; original draft preparation, G.L., A.S.; writing: G.L, A.S.; review and editing: B.R., A.D., F.L., M.P., F.V., S.M-A., G.L., A.S.; supervision: G.L., A.S.; funding acquisition: S.M-A., G.L., A.S.; project administration: G.L., A.S.; all authors have read and agreed to the published version of the manuscript.

## Data availability

The authors declare that data supporting the findings of this study are available within the article and its supplementary information files

## Declarations

### Ethics declarations

Informed Consent Statement: written informed consent was obtained from each patient involved in the study in compliance with the Helsinki declaration. Ethical standards set out in the Helsinki declaration were respected. The study was approved by the local bioethical committee of the University Hospital Center (Tours, France) and tissue and BALF collections were declared to the French Ministry of Higher Education, Research, and Innovation (approvals DC-2008-308, DC-2010-1216, and 2020/89196.1).

