## Supplementary figures for "Legumain (asparaginyl endopeptidase) modulates extracellular matrix dynamics in pulmonary fibrosis"

### ABSTRACT

Pulmonary fibrosis is characterized by extracellular matrix (ECM) deposition driven by fibroblast-to-myofibroblast transition (FMT) and by an altered proteolytic balance. While the roles of several cysteine proteases have been documented, the specific contribution of cathepsin V (CatV) and legumain (LGMN) remains poorly explored. LGMN, CatV and their dual inhibitor cystatin M/E (CysM/E) are significantly increased in lung specimens and bronchoalveolar lavage fluids from patients with idiopathic pulmonary fibrosis. TGF-β1 triggered CysM/E expression and LGMN transcription, intracellular maturation, enzymatic activity, and pro-LGMN secretion via the Smad-3 pathway, whereas CatV was downregulated in human lung fibroblasts (CCD-19Lu and primary HPF cells) undergoing myodifferentiation. Genetic silencing of LGMN or CatV, and pharmacological inhibition of LGMN, led to accumulation of fibronectin and elastin, implying that both proteases contribute to ECM remodeling. LGMN cleaved fibronectin, while CatV predominantly regulated elastin levels. Conversely, broad-spectrum inhibitor cystatin C (hCC) markedly reduced elastin and fibronectin degradation, whereas CysM/E exerted a weaker effect, mainly on elastin turnover. LGMN inhibition transiently delayed fibroblast wound closure, establishing a functional role in tissue repair through fibronectin remodeling. Neither LGMN nor CatV influenced α-SMA expression, distinguishing them from CatB, which participates in FMT. Altogether, LGMN was identified as an effector of matrix remodeling rather than myodifferentiation, acting in concert with CatV. Within the proteolytic network governing fibrosis progression, the present findings identify a cystatin-regulated LGMN/CatV partnership, participating in ECM turnover and cell migration. Present results also provide new perspectives on potential therapeutic protease-based strategies targeting ECM turnover underlying lung fibrosis.

**Keywords:** Cathepsin; Cystatin; Cysteine protease; ECM remodeling; Lung fibrosis; Wound healing.

a)

| Protein (Gene) | Strand (5' → 3') | Sequences of PCR primers |
|---|---|---|
| Ribosomal Protein S16 (RPS16) | Sense | ACGTGGCCCAGATTTATGCTAT |
| | Antisense | TGGAAGCCTCATCCACATATTTC |
| α-SMA (ACTA2) | Sense | CAGGGCTGTTTTCCCATCCAT |
| | Antisense | GCCATGTTCTATCGGGTACTTC |
| Fibronectin (FN1) | Sense | CGGTGGCTGTCAGTCAAAG |
| | Antisense | AAACCTCGGCTTCCTCCATAA |
| Elastin (ELN) | Sense | AACCAGCCTTGCCCCGC |
| | Antisense | CCCCAAGCTGCCTGGTG |
| Collagen I (COL1A1) | Sense | GTGCGATGACGTGATCTGTGA |
| | Antisense | CGGTGGTTTCTTGGTCGGT |
| Legumain (LGMN) | Sense | GCAGGTTCAAATGGCTGGTAT |
| | Antisense | GGAGTGGGATTGTCTTCAGAGT |
| Cathepsin V (CTSV) | Sense | ACAATGGGGAATACAGCCAAG |
| | Antisense | CAGAGGCTCACGGAACACTTT |
| Cystatin M (CST6) | Sense | TCCGAGACACGCACATCATC |
| | Antisense | CCATCTCCATCGTCAGGAAGTAC |
| Cystatin C (CST3) | Sense | GATCGTAGCTGGGGTGAACT |
| | Antisense | CCTTTTCAGATGTGGCTGGT |

b)

| Protein (Gene) | Name | siRNA target sequences |
|---|---|---|
| Legumain (LGMN) | Hs_LGMN_8 | GACGTGGAAGATCTGACTAAA |
| | Hs_LGMN_7 | CTGCCGGATAACATCAATGTT |
| | Hs_LGMN_6 | CGCAATGGGATTCCTGACGAA |
| | Hs_LGMN_3 | AAGGGTCATATTTGCTTTCTA |
| Cathepsin V (CTSV) | Hs_CTSL2_5 | TTCGTCCTTCCAGTTCTACAA |
| | Hs_CTSL2_4 | AAGACTCATTGCTTAATTCTA |
| | Hs_CTSL2_3 | CAGCAAGTATTGGCTCGTCAA |
| | Hs_CTSL2_1 | AAGGAGCAAATTCGAATAACA |
| Cystatin M (CST6) | Hs_CST6_8 | CAGCATCTACTACTTCCGAGA |
| | Hs_CST6_5 | CAAGTACTTCCTGACGATGGA |
| | Hs_CST6_3 | CAGGAGCGCATGGTAGGAGAA |
| | Hs_CST6_1 | CCGAGACACGCACATCATCAA |

**Supplementary Figure 1. Primer and small interfering RNA (siRNA) sequences.**
(**a**) Primers sequences used for real-time quantitative PCR (RT-qPCR). (**b**) Small interfering RNA (siRNA) sequences used in CDD-19Lu cells. hCC and scrambled siRNA sequences are the same as previously reported (Kasabova *et al.*, 2014a).

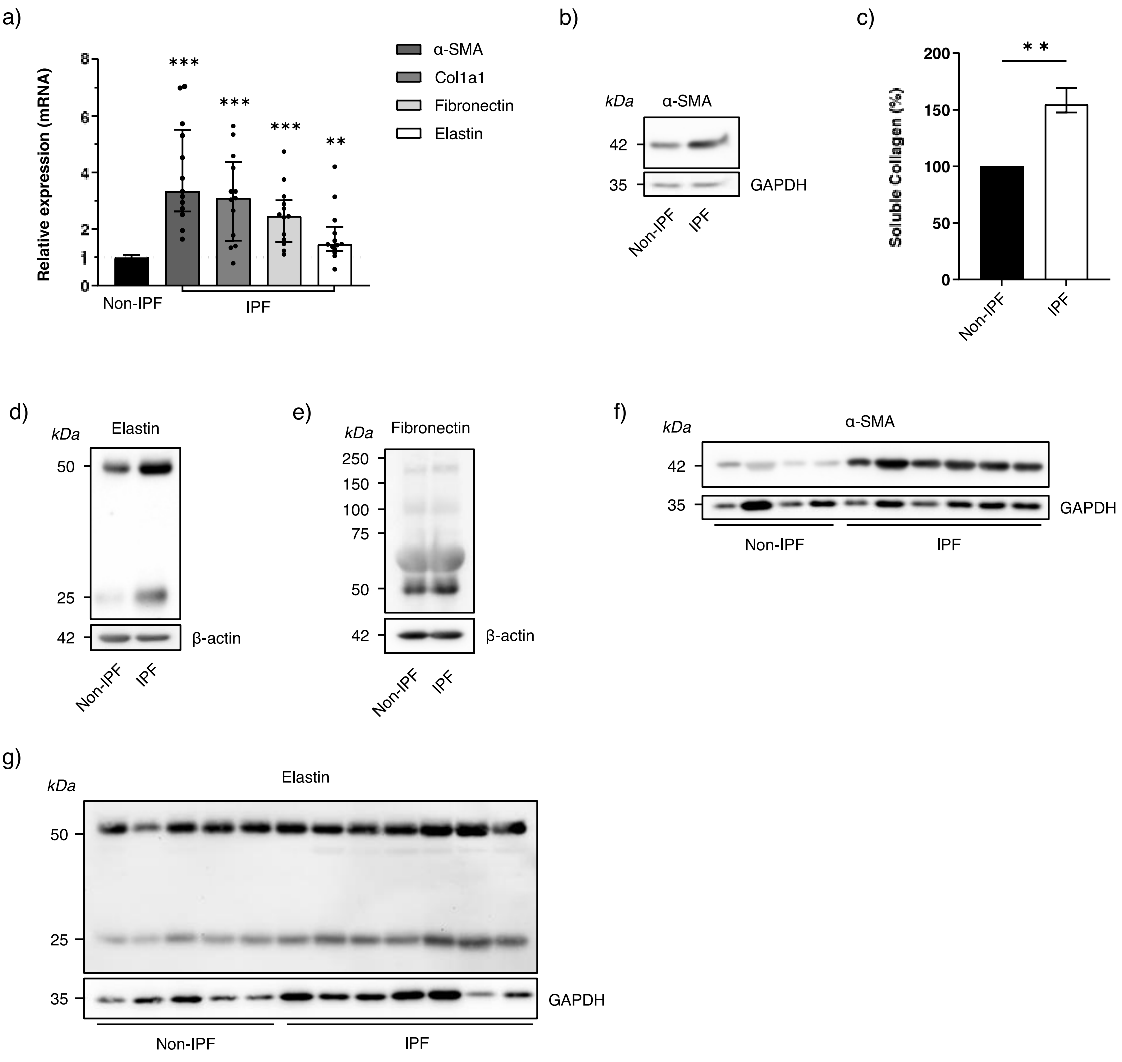


**Supplementary Figure 2. Expression level of pro-fibrotic markers in non-IPF and IPF human biopsies.**

(**a**) Transcriptional levels of α-SMA, Collagen I, fibronectin and elastin in human lung biopsies were analyzed by RT-qPCR. Data are expressed as relative fold change in non-IPF compared with IPF samples. (**b**) Western blot analysis of α-SMA in pooled lung biopsy extracts from non-IPF and IPF samples (100 µg protein, GAPDH as a loading control). (**c**) Quantification of soluble collagen in pooled lung biopsy extracts (Sircol collagen assay, n=4). The experiment was performed three times in duplicate. Western blot analysis of (**d**) elastin and (**e**) fibronectin in pooled lung biopsy extracts (50 µg protein, β-actin as a loading control). Western blot analysis of (**f**) and (**g**) α-SMA and elastin, respectively, in individual lung biopsy extracts (100 µg protein, GAPDH as a loading control). Bars represent median ± quartile. Statistical significance was assessed using the one-sample Wilcoxon test for RT-qPCR (** $p<0.01$; *** $p<0.001$) and the Mann-Whitney test for quantification of soluble collagen (** $p<0.01$).

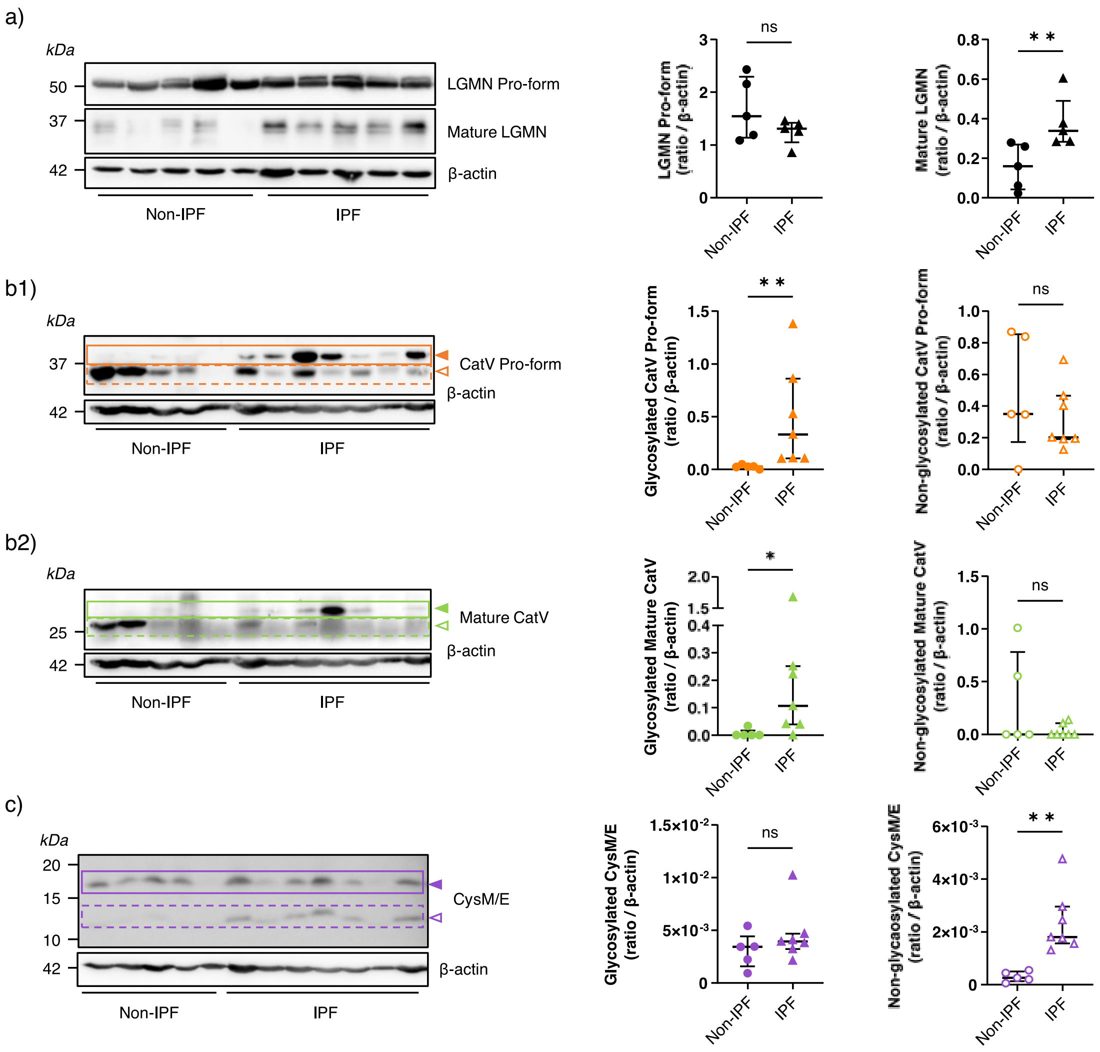


**Supplementary Figure 3. Expression levels of LGMN, CatV and CysM/E in non-IPF and IPF human biopsies.**

Western blot analysis of (**a**) LGMN, (**b.1**) CatV zymogen, (**b.2**) mature CatV, and (**c**) CysM/E in individual lung biopsy extracts from non-IPF and IPF samples (100 µg protein, β-actin as a loading control). Filled arrows indicate glycosylated forms, and open arrows indicate non-glycosylated forms. Densitometric analysis of protein expression levels was performed using ImageJ software and normalized to β-actin expression. Bars represent median ± quartile. Statistical significance was assessed using the Mann–Whitney test (ns: no significant; (* $p<0.05$; ** $p<0.01$).

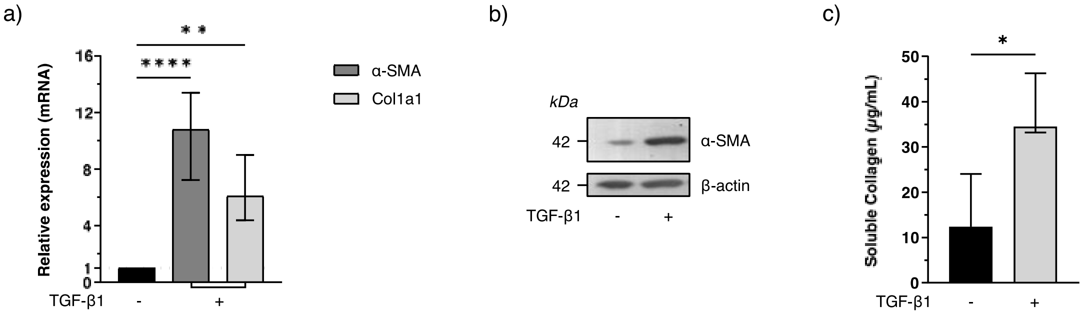


**Supplementary Figure 4. Effect of TGF-β1-dependant myodifferentiation on the expression of pro-fibrotic markers in human pulmonary fibroblasts (CCD-19Lu).**
CCD-19Lu cells were treated with recombinant human TGF-β1 (5 ng/mL) for 3 days to induce myofibroblast differentiation. (**a**) α-SMA and collagen I transcription levels were analyzed by RT-qPCR (n=5). Data are expressed as the relative fold change in myofibroblasts compared to fibroblasts. (**b**) α-SMA protein expression in cell lysates was analyzed by Western blot (20 µg protein, β-actin as a loading control). A representative blot from three independent experiments is shown. (**c**) Quantification of secreted soluble collagen (Sircol collagen assay, n=4). Bars represent median ± quartile. Statistical significance was assessed using the one-sample Wilcoxon test (* $p<0.05$, **** $p<0.0001$) for RT-qPCR data and the Mann-Whitney test (* $p<0.05$) for collagen quantification.

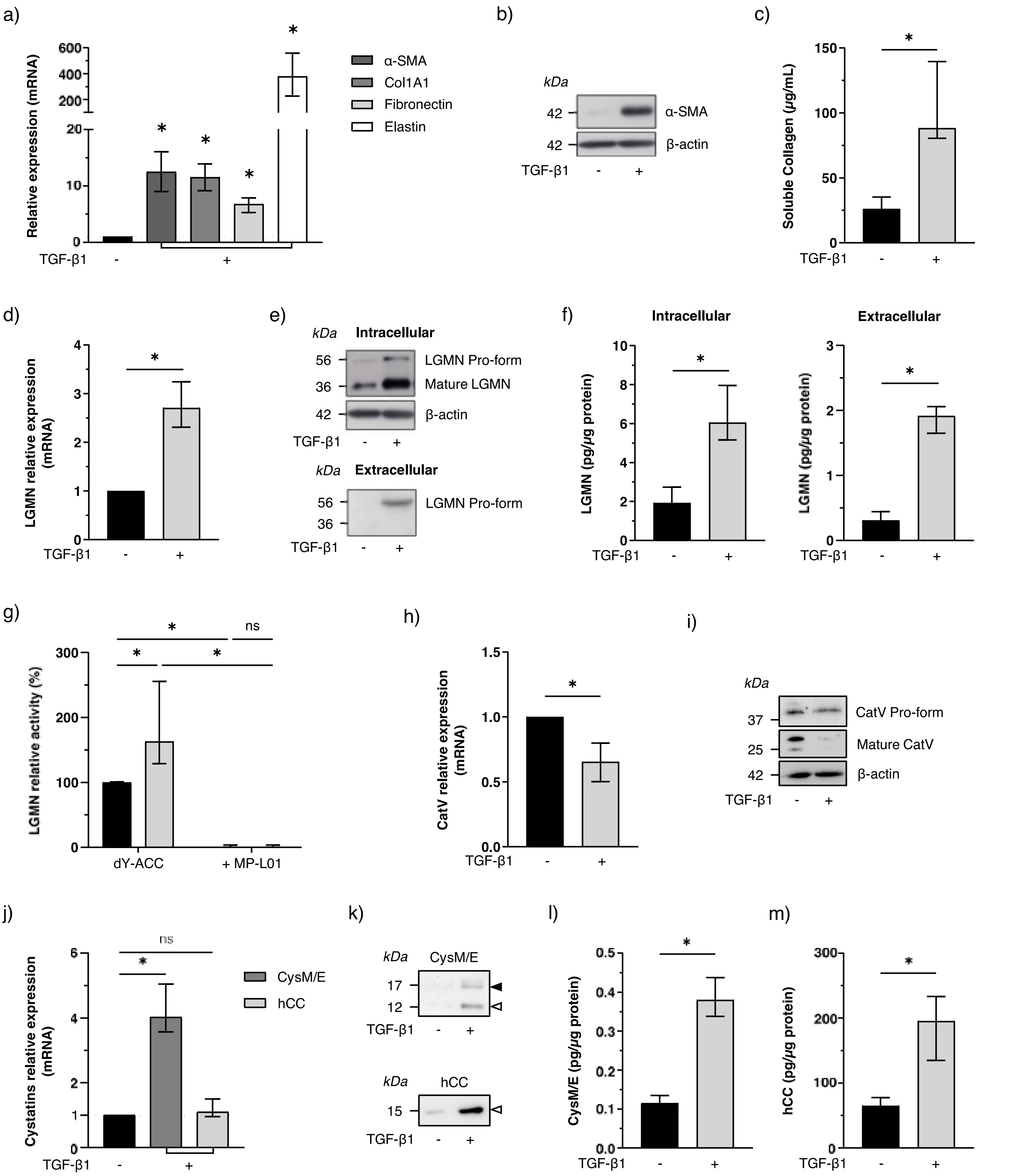


**Supplementary Figure 5. Effects of TGF-β1-dependant myodifferentiation on cysteine proteases, cystatins and pro-fibrotic markers in human pulmonary fibroblasts (HPF cells).**

HPF cells were treated with recombinant human TGF-β1 (5 ng/mL) for 3 days to induce myofibroblasts differentiation. (**a**) Transcription levels of α-SMA, collagen I, fibronectin and elastin were analyzed by RT-qPCR (n=4). (**b**) α-SMA protein expression in cell lysates was analyzed by Western blot (20 µg protein, β-actin as a loading control). (**c**) Quantification of soluble collagen secreted in culture medium (Sircol collagen assay, n=4). (**d**) LGMN transcription level was analyzed by RT-qPCR (n=4). (**e**) Western blot analysis of intracellular LGMN (30 µg protein, β-actin as a loading control) and secreted LGMN (50 µg protein). (**f**) ELISA quantification of total intracellular and secreted LGMN (n=4). (**g**) LGMN peptidase activity was measured in cell lysates (20 µg protein) using the fluorogenic substrate dY-ACC, with MP-L01 (10 µM) as a control inhibitor. Experiments were performed in duplicate (n=3). (**h**) CatV transcription level was analyzed by RT-qPCR (n=4). (**i**) CatV protein expression in cell lysates was analyzed by Western blot (100 µg proteins, β-actin as a loading control). (**j**)

Transcription levels of CysM/E and hCC were analyzed by RT-qPCR (n=4). (**k**) Western blot analysis of secreted CysM/E (100 µg protein) and hCC (50 µg protein) in culture medium. Black arrows indicate glycosylated forms, and white arrows indicate non-glycosylated forms. ELISA quantification of secreted (**l**) CysM/E and (**m**) hCC in culture medium (n=4). RT-qPCR data are expressed as the relative fold change in myofibroblasts compared with fibroblasts. A representative blot from three independent experiments is shown. Bars represent median ± quartile. Statistical significance was assessed using the Mann-Whitney test ( ns: not significant; * $p<0.05$) except for RT-qPCR data, which were analyzed using the one-sample Wilcoxon test (* $p<0.05$).

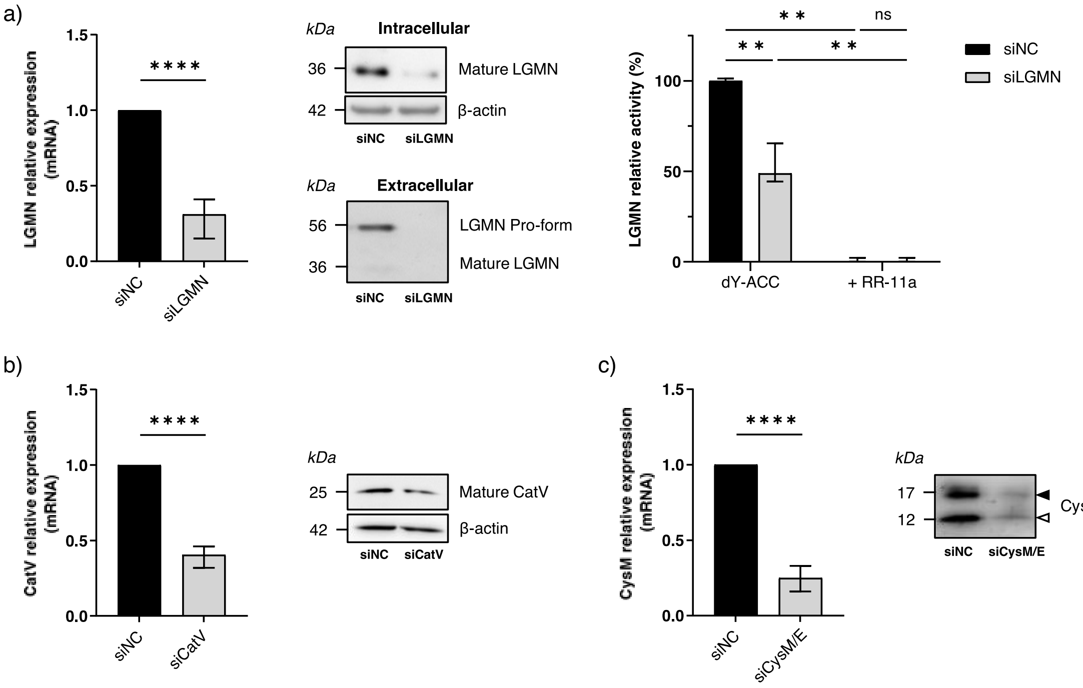


**Supplementary Figure 6. Validation of siRNA-mediated gene silencing in human lung myofibroblasts (CCD-19Lu).**

(**a**) LGMN expression after gene silencing was analyzed by RT-qPCR (n=5) and Western blot in cell lysates (30 µg protein, β-actin as a loading control) and culture medium (10 µg protein). LGMN peptidase activity was measured in cell lysates (20 µg protein) using fluorogenic substrate dY-ACC, with RR-11a (10 µM) as a control inhibitor. Experiments were performed in duplicate (n=5). Bars represent median ± quartile. (**b**) CatV expression after gene silencing was analyzed by RT-qPCR (n=5) and Western blot in cell lysates (50 µg protein, β-actin loading control). (**c**) CysM/E expression after gene silencing was analyzed by RT-qPCR (n=5) and Western blot in culture medium (10 µg protein). Black arrows indicate glycosylated forms, and white arrows indicate non-glycosylated forms. RT-qPCR data are expressed as relative fold change compared with scrambled siRNA-transfected cells. A representative blot from three independent experiments is shown. Statistical significance was assessed using the one-sample Wilcoxon test (**** $p<0.0001$) for RT-qPCR and the Mann-Whitney test (ns: not significant; ** $p<0.01$) for LGMN activity.

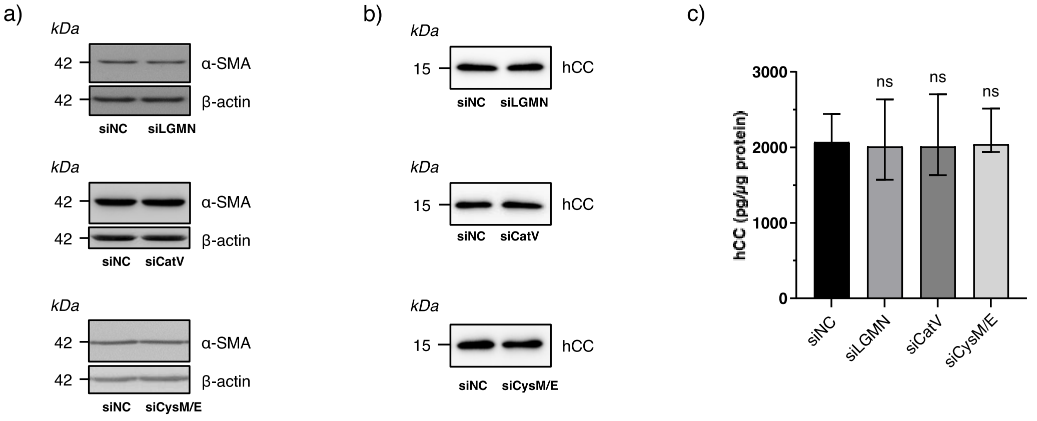


**Supplementary Figure 7. Effects of LGMN, CatV, and CysM/E silencing on myofibroblast differentiation markers (CCD-19Lu).**

(**a**) Western blot analysis of α-SMA protein expression in CCD-19Lu myofibroblasts transfected with LGMN, CatV, or CysM/E siRNAs compared with scrambled siRNA-transfected cells used as a negative control (20 µg protein, β-actin as a loading control). (**b**) Western blot analysis (5 µg protein) and (**c**) ELISA quantification (n=4) of hCC secreted by myofibroblasts transfected with LGMN, CatV, or CysM/E siRNA compared with scrambled siRNA-transfected cells used as a negative control. A representative blot from three independent experiments is shown. Bars represent the median ± quartile. Statistical significance was assessed using the Mann-Whitney test (ns: not significant).

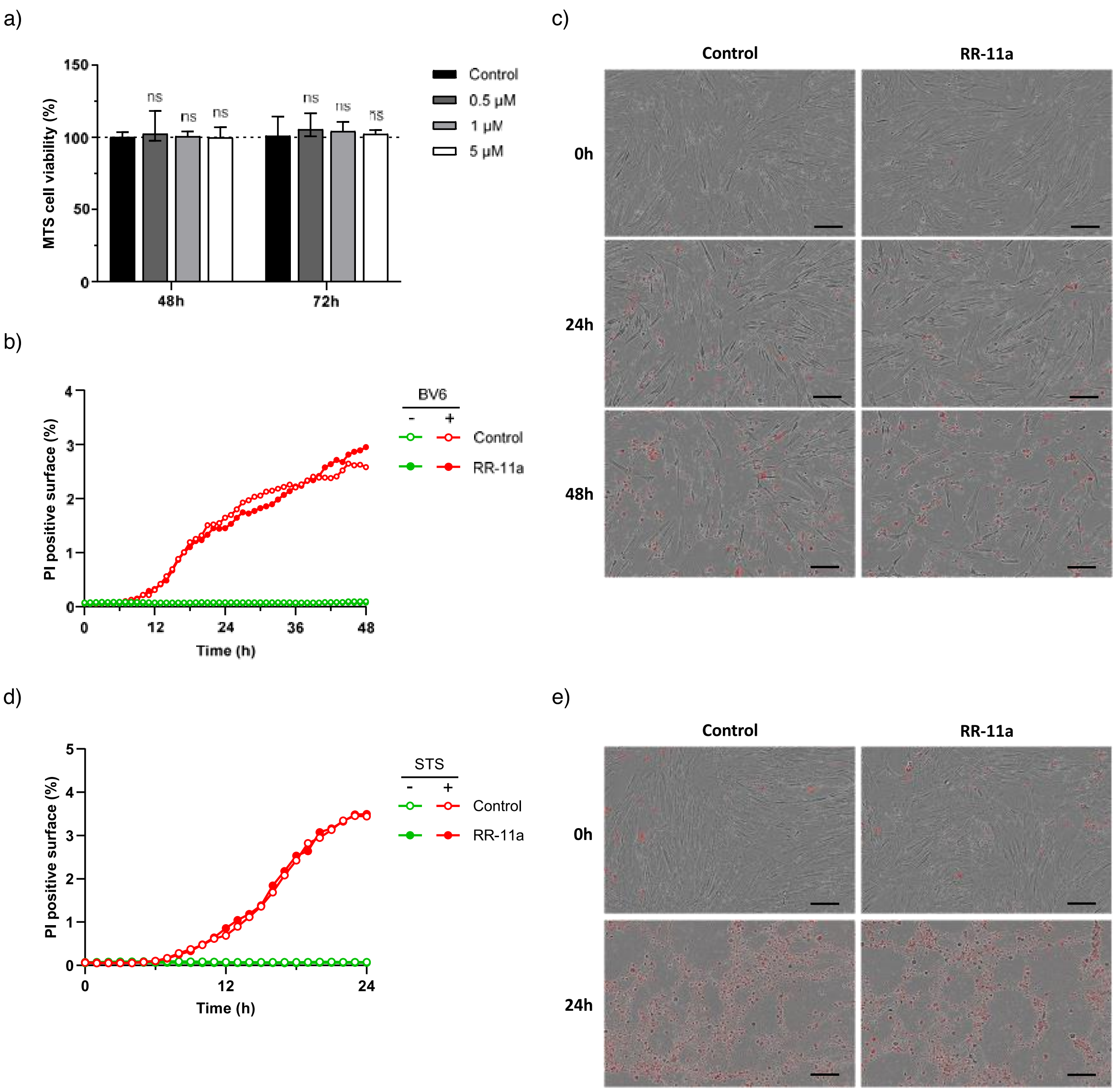


**Supplementary Figure 8. Effect of legumain inhibition on myofibroblasts viability and cell death.** (**a**) HPF cells were treated with recombinant human TGF-β1 (5 ng/mL) for 3 days to induce myofibroblast differentiation and were subsequently treated with RR-11a (0.5 – 5 µM) for 48 h to 72 h. Cell viability was assessed using the MTS assay. Bars represent the median ± quartile normalized to untreated cells (100% of viability) (n=4). (**b**) Myodifferentiated HPF cells were treated with BV6 (20 µM) to induce extrinsic cell death in presence or absence of RR-11a (1 µM). Cell death was monitored during 48 h and quantified with propidium iodide incorporation into the culture medium. Positive, red-stained cells due to PI were analyzed using IncuCyte software and results were expressed as the PI-positive area per image. (**c**) Representative images of BV6-induced cell death acquired at 0, 24 and 48 h (scale bar = 200 µm). (**d**) Myodifferentiated HPF cells were treated with staurosporine (STS; 0.5 µM) to induce intrinsic cell death in the presence or absence of RR-11a (1 µM). Cell death was monitored during 24 h and quantified with propidium iodide incorporation into the culture medium. (**e**) Representative images of STS-induced cell death acquired at 0 and 24 h (scale bar = 200 µm). Bars represent the median ± quartile. Statistical significance was assessed using the Mann-Whitney test (ns: not significant).
